# Regulation of RNA maturation by the family of human G-patch proteins

**DOI:** 10.64898/2026.06.30.735655

**Authors:** Indira Memet, Nidhi Kanwal, Anastasia Ritchie, Nicolai Krogh, Christof Lenz, A. Marieke Oudelaar, Henrik Nielsen, Henning Urlaub, Lydia Herzel, Katherine E. Bohnsack, Markus T. Bohnsack

## Abstract

The family of human G-patch proteins comprises more than 20 members, each characterized by a glycine-rich G-patch implicated in mediating interactions with RNA helicases. Here, we systematically identify the cognate RNA helicase of each G-patch protein, highlighting the association of DHX15 with a network of 20 G-patch cofactors. DHX35 and GPATCH1 represent a unique G-patch protein-RNA helicase pair, and we uncover a regulatory circuit between these partners. Comprehensive *in vitro* analyses of ATPase activity and RNA binding identify distinguishing features of DHX15- and non-DHX15-associated G-patch proteins, and demonstrate the roles of most G-patch proteins as *bona fide* stimulatory cofactors of DHX15. RNA interactome analyses of each G-patch protein and complementary transcriptome-wide alternative splicing analyses in cells lacking a G-patch protein reveal distinct modes of regulation of mRNA maturation by different G-patch proteins. For example, ZGPAT affects splicing indirectly through its requirement for efficient 2′-*O*-methylation of snRNAs, GPATCH8 exemplifies DHX15-associated alternative splicing modulation, whereas SUGP2 suppresses splicing in an RNA helicase-independent manner via direct binding to pre-mRNA introns.

## INTRODUCTION

Gene expression is a fundamental cellular pathway in which genomic information is transcribed to produce messenger RNA (mRNA) precursors that undergo extensive maturation before they are translated by ribosomes to synthesize the cellular proteome. Already co-transcriptionally, numerous proteins are recruited to pre-mRNAs and the protein interactomes of (pre-)mRNAs dynamically alter during their lifecycle^1^. Pre-mRNA biogenesis involves 5′ end capping, 3′ end polyadenylation, installation of internal RNA modifications, folding and association with RNA-binding proteins (RBPs) as well as the removal of introns by pre-mRNA splicing. Excision of introns is mediated by spliceosomal complexes that assemble *de novo* on each intron and enable the two *trans*esterification reactions that form the intron lariat and then close the exon-exon junction^2^. Pre-mRNA splicing is facilitated by small nuclear RNPs (snRNPs), composed of numerous proteins as well as small nuclear RNAs (snRNAs) that base-pair with specific regions of the pre-mRNA substrate, tethering them in appropriate conformations for the splicing reactions to occur^3,4^. Alternative pre-mRNA splicing, which can manifest as exon skipping, the inclusion of mutually exclusive exons, use of alternative 5′ or 3′ splice sites (ss) or retention of introns, substantially increases the diversity of the transcriptome and represents an efficient mechanism via which the proteome can be dynamically adapted in different cell types and environmental conditions^5^.

Among the plethora of RBPs that associate with pre-mRNAs during their maturation^6,7^ there are several examples of proteins that share common features endowing specific properties, such as enzymatic activities or the ability to mediate protein–protein or protein–RNA interactions. For example, serine-arginine-rich (SR) proteins are prominent components of spliceosomal complexes that are characterized by their ability to interact simultaneously with pre-mRNAs via an RNA recognition motif (RRM) and with protein components of the splicing machinery through an RS domain rich in arginine and serine residues^8^. Another element shared by numerous RBPs implicated in, among other cellular process, pre-mRNA splicing is the G-patch^9^.

The approximately 50 amino acid long, glycine-rich G-patch is present in >20 human proteins, and is characterized by seven highly conserved glycine residues and an invariable aromatic amino acid interspersed with three hydrophobic patches (consensus sequence: Gx_2_hhx_3_Gax_2_GxGlGx_3_pxux_3_sx_10-16_GhG where a = aromatic, h = hydrophobic, l = aliphatic, p = polar, s = small, u = tiny, x = variable amino acid; **Supplementary Fig. S1a**)^10^. G-patch-containing proteins are present throughout eukaryotes, and G-patches are also encoded within several beta-retroviruses and endogenous retroviral elements^11,12^. On the structural level, the G-patch comprises a short N-terminal a-helix followed by an unstructured region. While G-patches are only present in single copies within proteins, they are often accompanied by domains/motifs mediating protein–protein interactions and/or RNA/DNA binding (**Fig. 1a**).

**Figure 1.**
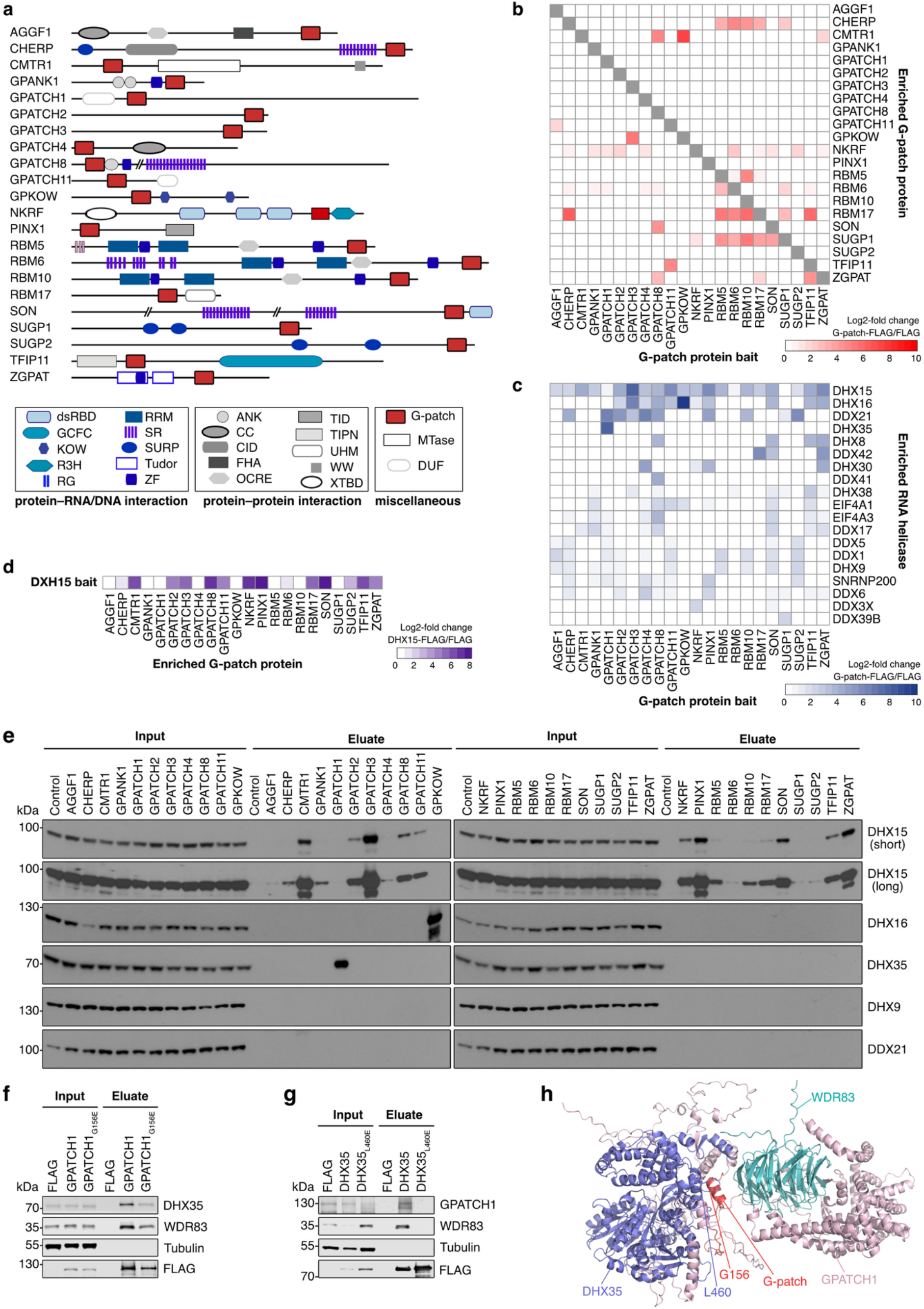
G-patch protein interactions with RNA helicases in cells. **(a)** Scheme of the domain and motif architecture of the human G-patch proteins. Proteins and elements within are drawn to scale with the exception of SON wherein // indicates 650 amino acids. Abbreviations: dsRBD – double-stranded RNA binding domain, GCFC – GC-rich sequence DNA-binding factor-like protein, HMG – high-mobility group box, KOW – Kyprides, Ouzounis, Woese motif, R3H – R3H motif, RG – arginine-glycine repeats, RRM – RNA recognition motif, SR – serine-arginine repeats, SURP – suppressor-of-white-apricot and PRP21/SPP91, Tudor – Tudor domain, ZF – Zinc finger, ANK – ankyrin repeat, CC – coiled coil, CID – RNA polymerase 2 C-terminal domain interacting domain, FHA – forkhead-associated domain, OCRE – octamer repeat domain, SOX – SOX17/18 central domain, TID – telomerase inhibitory domain, TIPN – Tuftelin interacting protein N-terminal domain, UHM – U2AF homology motif, WW – two tryptophan containing domain, XTBD – XRN2 binding domain, MTase – ribose 2’-*O*-methyltransferase, DUF – domain of unknown function. **(b, c)** Lysates from stably transfected cell lines expressing the FLAG tag or FLAG-tagged G-patch proteins were used for IP experiments and eluates were analyzed by MS. Heatmap showing the log2-fold enrichment of G-patch proteins compared to the FLAG control (b). Heatmap showing the log2-fold enrichment of RNA helicases with the G-patch protein baits compared to the FLAG control (c). **(d)** Lysates from stably transfected cell lines expressing the FLAG tag or FLAG-tagged DHX15 were used for IP experiments and elutes were analyzed by MS. Heatmap showing the log2-fold enrichment of G-patch proteins with DHX15 compared to the FLAG control. **(e)** IPs were performed as in (b, c), and inputs (1%) and eluates were analyzed by western blotting using antibodies against the indicated proteins. Two exposures (long and short) are shown for DHX15. **(f, g)** IPs were performed using lysates from stably transfected cell lines expressing the FLAG tag, or wild-type or mutant GPATCH1 (f) or DHX35 (g). Inputs (1%) and eluates were analyzed by western blotting using antibodies against the indicated proteins. **(h)** Model of DHX35–GPATCH1-WDR83 complex in which DHX35 is shown in slate, GPATCH1 in light pink and WDR83 in teal. The G-patch of GPATCH1 (152-198) is highlighted in red and the amino acids substituted are marked.

Structural and functional studies indicate that the G-patch motif can mediate interactions between G-patch proteins and RNA helicases of the DExH family (see for example^13–20^). RNA helicases are nucleoside triphosphate (NTP)-dependent enzymes that provide the driving force for decisive remodeling steps of RNAs and ribonucleoprotein complexes (RNPs) through their mechanical functions in rearranging RNA structures and displacing RNA-binding proteins^21^. RNA helicases are composed of tandem RecA-like domains tethered by a flexible linker and often flanked by variable N- and C-terminal domains. RNA helicases of the DExH sub-family are translocating unwindases; cycles of NTP hydrolysis trigger the RecA2 domain to toggle between open and closed conformations where each opening step is coupled to a one nucleotide (nt) movement of the RNA substrate through the RNA-binding channel of the helicase^22,23^. On the cellular level, precise regulation of RNA helicase activity is critical, but the helicase core predominantly contacts the RNA substrate backbone^24^, indicating a lack of inherent sequence specificity. Protein cofactors of RNA helicases often couple recruitment of the enzyme to appropriate substrates RNAs/RNPs with allosteric stimulation of its catalytic activity to ensure targeted RNA helicase action within the RNA-dense cellular environment^25^. Structural analyses of the G-patch motifs of NKRF (human) and Spp2 (*S. cerevisiae*) with their associated RNA helicases, DHX15 (NKRF) and Prp2 (Spp2) reveal that the G-patch motif binds in an extended conformation across the helicase surface; the N-terminal a-helix of the G-patch domain contacts the winged-helix region of the helicase while the unstructured C-terminal region extends into a pocket within the RecA2 domain^19,20^. In the case of NKRF, this arrangement tethers the catalytic core of the cognate helicase in an optimal confirmation for RNA binding and NTP hydrolysis, thus rationalizing the stimulatory effect of the G-patch protein on the helicase^16,20^.

The mosaic architecture of G-patch proteins, especially the common co-occurrence of G-patch motifs alongside nucleic acid binding domains/motifs suggests that these proteins typically function within ribonucleoprotein complexes (RNPs) in the context of gene expression^9^. Much functional knowledge on the regulation of RNA helicases by G-patch proteins comes from the *S. cerevisiae* model system. Four yeast G-patch proteins stimulate the activity of the DExH box protein Prp43 with functions for these helicase–G-patch protein complexes described in ribosome assembly and pre-mRNA splicing^26–29^. Interestingly, alterations in the levels of these Prp43-associated G-patch proteins regulate the distribution of this multifunctional helicase between its different cellular functions^27^. The yeast G-patch protein, Spp2, acts together with its associated DExH protein Prp2 to mediate formation of catalytically active spliceosomes^30^. The yeast G-patch proteins are conserved in humans, but the human G-patch protein family includes numerous additional members^9^. Some human G-patch proteins have been linked to important and diverse cellular functions. For example, CMTR1 2′-*O*-methylates the cap-proximal nucleotide of pre-mRNAs^31,32^, ZGPAT is a transcriptional repressor^33^, PINX1 functions as a telomerase inhibitor^34^, SON nucleates formation of nuclear speckles^35^, AGGF1 is a well-characterized angiogenesis factor^36^ and RBM5 acts as a potent tumor suppressor^37^. Several G-patch proteins have also been shown to associate with the multifunctional RNA helicase, DHX15, the human homologue of Prp43^3–17,28,38–40,41^. Despite these findings, many human G-patch proteins remain functionally uncharacterized and often, the role of the G-patch motif in ascribed cellular functions is unclear.

Here, we systematically analyze the human G-patch protein family, exploring their roles as RNA helicase cofactors and sub-cellular localizations, as well as their RNA interactomes and influence on alternative pre-mRNA splicing.

## RESULTS

### Defining the RNA helicase interactome of the human G-patch proteins

The human G-patch proteins represent a family of putative RNA helicase cofactors but for various G-patch proteins, an RNA helicase interaction partner remains unknown. To systematically identify cognate helicases of the human G-patch proteins, stably transfected HEK293 cell lines for the inducible expression of C-terminally FLAG-tagged versions of each of these proteins were generated (**Supplementary Fig. S1b**). These cell lines, together with a control cell line expressing only the FLAG tag, were then used for immunoprecipitation (IP) experiments, performed in the presence of RNase, and co-precipitated proteins were detected by mass spectrometry (MS). Comparison of the proteomes of the IP eluates from cells expressing FLAG-tagged G-patch proteins with that of the FLAG control enabled the identification of G-patch protein-associated proteins (**Supplementary Data 1**). Each of the G-patch protein baits was efficiently recovered and interestingly, several G-patch proteins also recovered other G-patch proteins (**Fig. 1b**). For example, RBM10 enriched RBM5, and RBM5 itself substantially recovered RBM6, findings which are in line with previous observations that these proteins functionally cooperate^42,43^. Similarly, CHERP and RBM17 reciprocally recovered each other, consistent with their reported co-functionality^44^. Furthermore, strong associations were observed between the RBM5/RBM6/RBM10 cluster and CHERP/RBM17, and SUGP1 was also found clearly to enrich, or be enriched by, all these G-patch proteins (**Fig. 1b**). GPKOW strongly enriched the cap-proximal mRNA 2′-*O*-methyltransferase CMTR1 and was itself recovered with GPATCH3, whereas GPATCH8 co-purified SON, CMTR1 and ZGPAT. TFIP11, likewise, networks with several other G-patch proteins; both RBM17 and ZGPAT were enriched with TFIP11, and TFIP11 was recovered with GPATCH11 (**Fig. 1b**). Together, these data suggest that while some G-patch proteins exert their functions independently of other G-patch proteins, others exist within macromolecular complexes containing multiple G-patch proteins.

Focusing then on RNA helicases, revealed DHX15 to be the most enriched DExH helicase for 19 human G-patch proteins (**Fig. 1c**). Consistent with this, 14 of these G-patch proteins were also co-precipitated in an IP experiment performed using FLAG-DHX15 as a bait (**Fig. 1d**; **Supplementary Data 2**). Among the G-patch proteins enriched with DHX15 was RBM6, which itself recovered DHX15 only weakly, but SUGP1, which lacked a DExH helicase interactor in its IP-MS was not significantly co-purified. Western blotting of IP eluates confirmed the interactions of all 19 G-patch proteins with DHX15 (**Fig. 1e**) and demonstrated a weak, but reproducible interaction with SUGP1, as recently reported^45^. The DEAD box ATPase DDX21, which was identified as enriched in several G-patch protein IP eluates by MS, was not confirmed as a *bona fide* interactor by the western blotting analysis, implying that it is a frequent contaminant highlighted by MS. As expected by analogy to yeast and consistent with previous data^46^, only GPKOW associated with DHX16 more strongly than DHX15 (**Supplementary Data 1**; **Fig. 1c, e**). While DHX16 was present in the interactomes of various other G-patch proteins, its recovery was less than DHX15 (**Fig. 1c**) and it was below the detected threshold of western blotting (**Fig. 1e**), suggesting that its co-precipitation reflects its presence in larger assemblies containing G-patch protein–DHX15 complexes. The remaining G-patch protein, GPATCH1, co-purified neither DHX15 nor DHX16, but instead DHX35 was specifically recovered (**Supplementary Data 1**; **Fig. 1c**) and also confirmed to associate (**Fig. 1e**). Analysis of helicase–G-patch interface residues across the DExH family members suggested that only DHX15, DHX16 and DHX35 could bind G-patch proteins due to the presence of disruptive amino acids in all other helicases^20^, and our data demonstrate that each of these helicases does indeed associate with a, or multiple, G-patch protein(s) in cells. Despite apparently sharing a common interaction interface, each G-patch protein appears to associate with a specific cognate DExH helicase (**Fig. 1e**). Although the IPs were performed under stringent conditions with the aim of predominantly enriching direct interaction partners of the G-patch proteins, the inventories of recovered proteins (**Supplementary Data 1**) also provide insights into the cellular contexts of these proteins (see below). Together these results comprehensively identify DExH helicase interaction partners of all human G-patch proteins, revealing that DHX15 associates with an extensive network of 20 cofactor proteins, and uncovering DHX35 and GPATCH1 as a distinct DExH/RHA helicase-G-patch protein partnership in human cells.

Structural analyses of DHX15 and the G-patch (GP) of NKRF^20^, and yeast Prp2 (DHX16 in humans) and the G-patch of Spp2^19^, demonstrated that N-terminal a-helices (brace helix) of the G-patches contact the winged-helix (WH) domains of the helicases while the C-terminal loop (brace loop) of the G-patches interact with the helicase RecA2 domains. To determine whether DHX35 and GPATCH1 interact in an analogous manner, IP experiments were performed using extracts from cell lines expressing FLAG-tagged wild-type GPATCH1/DHX35 or versions containing amino acid substitutions, which by analogy to the DHX15-NKRF_GP_ structure^20^ should perturb G-patch protein–helicase interaction. FLAG-tagged GPATCH1 in which glycine 156 within the brace helix was substituted with glutamate (GPATCH1_G156E_) was expressed as well as FLAG-tagged DHX35 containing a leucine to glutamate substitution at position 460 within the putative G-patch brace helix interaction interface in the winged helix domain (DHX35_L460E_). Wild-type GPATCH1 robustly recovered DHX35 and reciprocally, wild-type DHX35 retrieved GPATCH1 (**Fig. 1f, g**). However, both the GPATCH1_G156E_ and DHX35_L460E_ substitutions diminished these interactions (**Fig. 1f, g**), implying that GPATCH1 and DHX35 interact analogously to other DExH–G-patch proteins. In the yeasts *S. pombe* and *C. thermophilum*, Gpatch1 and Dhx35 associate with Wdr83^47,48^, and alongside DHX35, human WDR83 was found to be highly enriched in the IP analyses of GPATCH1 (**Supplementary Data 1**). Robust interactions between WDR83 and both GPATCH1 and DHX35 were also demonstrated (**Fig. 1f, g**), indicating that this trimeric complex also forms in human cells. Detection of WDR83 in the IP experiments revealed that although the G156E substitution in GPATCH1 affected its interaction with DHX35, the amount of WDR83 recovered with GPATCH1 was unaffected (**Fig. 1f**). By contrast, the L460E substitution in DHX35 impaired its interaction not only with GPATCH1 but also WDR83 (**Fig. 1g**). Together, these results indicate a DHX35-independent association between GPATCH1 and WDR83, and that GPATCH1 bridges the interaction between WDR83 and DHX35. This notion is supported by modeling of the DHX35–GPATCH1–WDR83 complex (**Fig. 1h**; **Supplementary Fig. S1c**); the compact WD repeat-containing protein WDR83 is placed such to form extensive interactions with regions of GPATCH1 outside the G-patch domain, which itself is positioned proximal to the WH/RecA2 domains of DHX35.

### Regulation of DExH box helicase activities by G-patch proteins

*Bona fide* RNA helicase cofactors facilitate recruitment to appropriate substrates and/or act as positive or negative regulators of the catalytic activity of their cognate helicases^25^. To determine whether all G-patch protein–DExH helicase interactions identified represent functional interactions that regulate helicase activity, the influence of each G-patch on RNA binding and ATP hydrolysis by the cognate helicases was analyzed. For this, MBP-/His_10_-tagged DHX15, DHX16 and DHX35, and ZZ-/His_7_-tagged versions of the human G-patches (G-patch protein_GP_), as well as the ZZ-His_7_ tag alone were recombinantly expressed in *Escherichia coli* (*E. coli*) and purified by affinity chromatography (**Fig. 2a-c**). The steady-state ATP hydrolysis activity of DHX15 or DHX16 in the presence of RNA and either the ZZ-His_7_ tag or each of the ZZ-His_7_ tagged G-patch domains was monitored using an *in vitro* NADH-coupled ATPase assay (**Fig. 2d; Supplementary Data 3**). The G-patch domains of all the DHX15-interacting G-patch proteins, which do not themselves hydrolyze ATP (**Supplementary Data 3**), stimulated the ATPase activity of DHX15 but not DHX16, with the exception of the RBM6_GP_, which did not influence the activity of either helicase (**Fig. 2e**). RBM6 was one of the G-patch proteins for which DHX15 was most weakly recovered by IP-MS (**Fig. 1e**), suggesting that the interaction between these proteins is weak or transient, which could rationalize the failure of this G-patch to stimulate the helicase *in vitro*, despite the presence of the key amino acids of the G-patch motif (**Supplementary Fig. S1a**).

**Figure 2.**
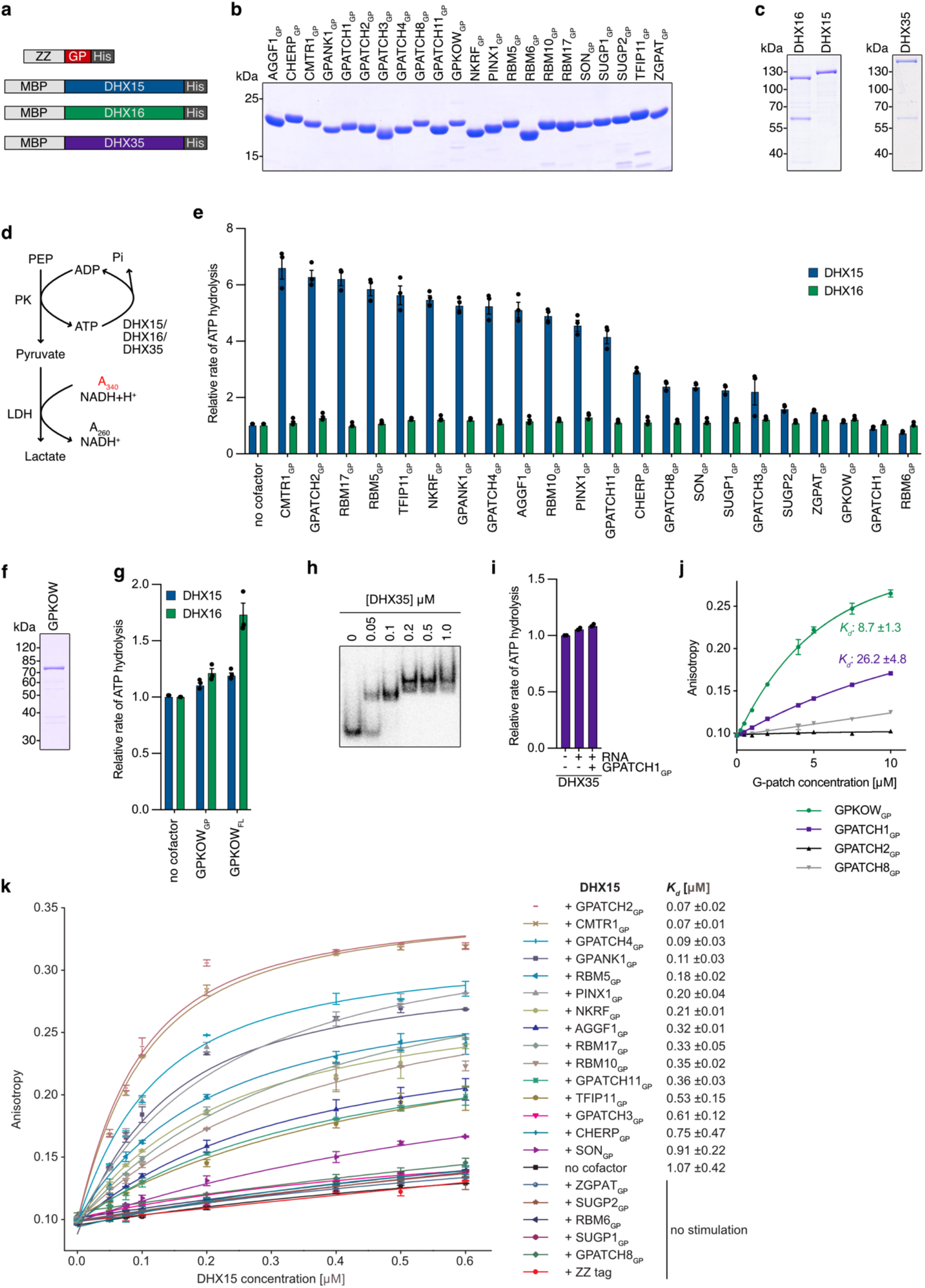
G-patch protein interactions with RNA helicases in cells. **(a)** Schemes of the recombinant proteins used. **(b, c)** N-terminally ZZ- and C-terminally His_7_-tagged versions of the G-patches (see Supplementary Fig. 1a) of the human G-patch proteins (b), and N-terminally MBP- and C-terminally His_10_-tagged versions of DHX15, DHX16 and DHX35 (c) were recombinantly expressed in *E. coli* and purified by affinity chromatography. Purified proteins were separated by SDS-PAGE and visualized by Coomassie staining. **(d)** Scheme of the NADH-coupled ATPase activity assay. Conditions were titrated such that ATP hydrolysis by the RNA helicase is the rate-limiting step. **(e)** The relative rates of ATP hydrolysis by DHX15 and DHX16 (250 nM) in the presence of RNA but without cofactor (no G-patch), and in the presence of both RNA (U_32_; 2 μM) and G-patches (1.5 μM), were determined. Data are presented as mean of n=3 experiments ± standard error of the mean (sem). **(f)** N-terminally ZZ- and C-terminally His_7_-tagged full-length GPKOW (GPKOW_FL_) was recombinantly expressed in *E. coli* and purified by affinity chromatography. Purified protein was separated by SDS-PAGE and visualized by Coomassie staining. **(g)** The relative rates of ATP hydrolysis by DHX16 (250 nM) in the presence of RNA (U_32_; 2 μM) with and without GPKOW_GP_ or GPKOW_FL_ (1.5 μM) were determined. Data are presented as mean of n=3 experiments ± standard error of the mean (sem). **(h)** Binding of MBP-DHX35-His10 to a 5’ [^32^P]-labelled U_32_ RNA was analyzed by electromobility shift assay. RNA migration in the native polyacrylamide gel was detected using a phosphorimager. **(i)** The relative rates of ATP hydrolysis by DHX35 (250 nM) in the absence or presence of RNA (U_32_; 2 μM) and with RNA and GPATCH1_GP_ (1.5 μM) were determined. Data are presented as mean of n=2 experiments ± standard error of the mean (sem). **(j)** Fluorescence anisotropy measurements were performed using an ATTO647N-labelled U_32_ RNA and different concentrations (0-10 μM) of the indicated G-patches. Results from n=3 experiments are shown as mean± sem. The dissociation constants (*K*_*d*_) are given for GPKOW_GP_ and GPATCH1_GP_, and were in the mM range for GPATCH2_GP_ and GPATCH8_GP_. **(k)** Fluorescence anisotropy measurements were performed using an ATTO647N-labelled U_32_ RNA un the presence of increasing amounts of DHX15 (0-0.6 μM) either without cofactor or with addition of 1.2 μM of the indicated ZZ-/His_7_-tagged G-patches or the ZZ-/His_7_ tag only. Data from n=2 experiments are shown as mean± standard error and the dissociation constants (*K*_*d*_) are given.

Only a very mild stimulation of DHX16 by the G-patch of its associated G-patch protein GPKOW was observed (**Fig. 2e**). Even in the absence of spliceosomes, the yeast homologue of GPKOW, Spp2, promotes ATP hydrolysis by Prp2 (yeast DHX16)^30^, so to determine if regions of GPKOW outside the G-patch contribute to DHX16 interaction and ATPase regulation, full-length GPKOW was recombinantly expressed and purified (**Fig. 2f**). Although still less than 2-fold, full-length GPKOW stimulated the ATPase activity of DHX16 more strongly than the GPKOW G-patch alone (**Fig. 2g**), suggesting that this is indeed the case and that, the G-patch protein has the potential to promote ATP hydrolysis by the helicase, albeit weakly outside the spliceosomal context. To date, DHX35 has not been demonstrated to be an active ATPase so the influence of RNA as well as the G-patch of GPATCH1 was explored. DHX35 alone hydrolyzed ATP poorly and, although DHX35 bound to the RNA substrate (**Fig. 2h**), this did not stimulate ATPase activity, neither alone nor in the presence of GPATCH1_GP_ (**Fig. 2i**).

Structural analyses indicate that G-patches can act as flexible linkers, tethering the two RecA-like domains of the helicase in closer proximity^19,20^. For several RNA helicases, co-operativity between RNA substrate binding and ATPase activity has been observed^21^, raising the possibility that the extents of ATPase activation of DHX15, DHX16 and DHX35 observed correspond to influences of the various G-patches on RNA affinities of the helicases. Prior to investigating whether the G-patches influence RNA binding by their cognate RNA helicases, it was necessary to ascertain whether the G-patches themselves bind to RNA. Strikingly, fluorescence anisotropy experiments revealed that while the DHX15-interacting G-patches do not stably interact with RNA (**Supplementary Data 4**), the G-patches of both GPKOW and GPATCH1, which interact with DHX16 and DHX35, respectively, bound the RNA substrate with *K*_*d*_ values in the low mM range (**Fig. 2j**). This suggests that features of the GPATCH1 and GPKOW G-patches that render their interactions with DExH box helicases specific for DHX16 and DHX35 may endow RNA binding ability or co-exist with elements that do so.

As GPATCH1_GP_ and GPKOW_GP_ bound RNA, their influences on the RNA affinities of DHX35 and DHX16 were not explored. However, fluorescence anisotropy experiments were conducted for DHX15 in the absence and presence of each of its associated G-patches (**Fig. 2k**). Under the conditions used, in the absence of any G-patch or in the presence of the ZZ-His_7_ tag, DHX15 bound the RNA substrate with an estimated *K*_*d*_ of approximately 1 μM. Addition of most individual G-patch domains stimulated RNA binding by DHX15 with affinities reaching up to *K*_*d*_ 70 nM (GPATCH2_GP_, CMTR1_GP_) (**Fig. 2k**). The extent of enhancement of RNA binding by DHX15 correlated well with the extent of stimulation of ATPase activity, and only the G-patches that minimally stimulated ATP hydrolysis by DHX15 did not enhance RNA binding by the helicase (**Fig. 2e, k**).

Together, these results indicate that the G-patch is sufficient to confer specificity for the appropriate helicase. The G-patches of DHX16/DHX35-interacting G-patch proteins do not, or only very weakly, influence ATPase activity of their associated helicases, whereas almost all DHX15-interacting G-patches promote RNA binding and consequently ATP hydrolysis by the helicase. In contrast to the DHX15-interacting G-patches, GPKOW_GP_ and GPATCH1_GP_ have the capacity to bind RNA themselves. Dissecting the regulatory effects of GPKOW and GPATCH1 on their cognate helicases requires further dedicated efforts, but the DHX15-interacting G-patch proteins can be collectively considered a network of stimulatory helicase cofactors.

### Human G-patch proteins localize within the nucleoplasm and to various sub-nuclear compartments

Insights into the processes in which particular proteins act can often be gained from their sub-cellular localization. Therefore, immunofluorescence microscopy was performed on the cell lines using an anti-FLAG antibody to detect the G-patch protein. Nuclear material was visualized by DAPI staining and immuno-detection of the well characterized ribosome assembly factor UTP14A ^49^ served as a marker for nucleoli (**Fig. 3**). Four G-patch proteins were detected in nucleoli: NKRF and GPATCH4 exclusively, and GPATCH2 and PINX1 together with a nucleoplasmic signal. The detection of NKRF and GPATCH4 in nucleoli is consistent with the co-precipitation of various ribosomes assembly factors (**Supplementary Data 1**; **Fig. 4a**) and their recently described roles in pre-rRNA processing and facilitating efficient rRNA modification^16,17^. GPATCH2 and PINX1 are the homologues of the yeast G-patch proteins Sqs1 (alias Pfa1) and Prx1 (alias Gno1)^28,50^, which are also implicated in ribosome assembly^28,29,51^. The localization of GPATCH2/PINX1 to nucleoli in human cells, taken together with the enrichment of numerous ribosome assembly factors in the IP-MS analyses of these proteins (**Supplementary Data 1**; **Fig. 4a**), suggests that their functions are likely conserved.

**Figure 3.**
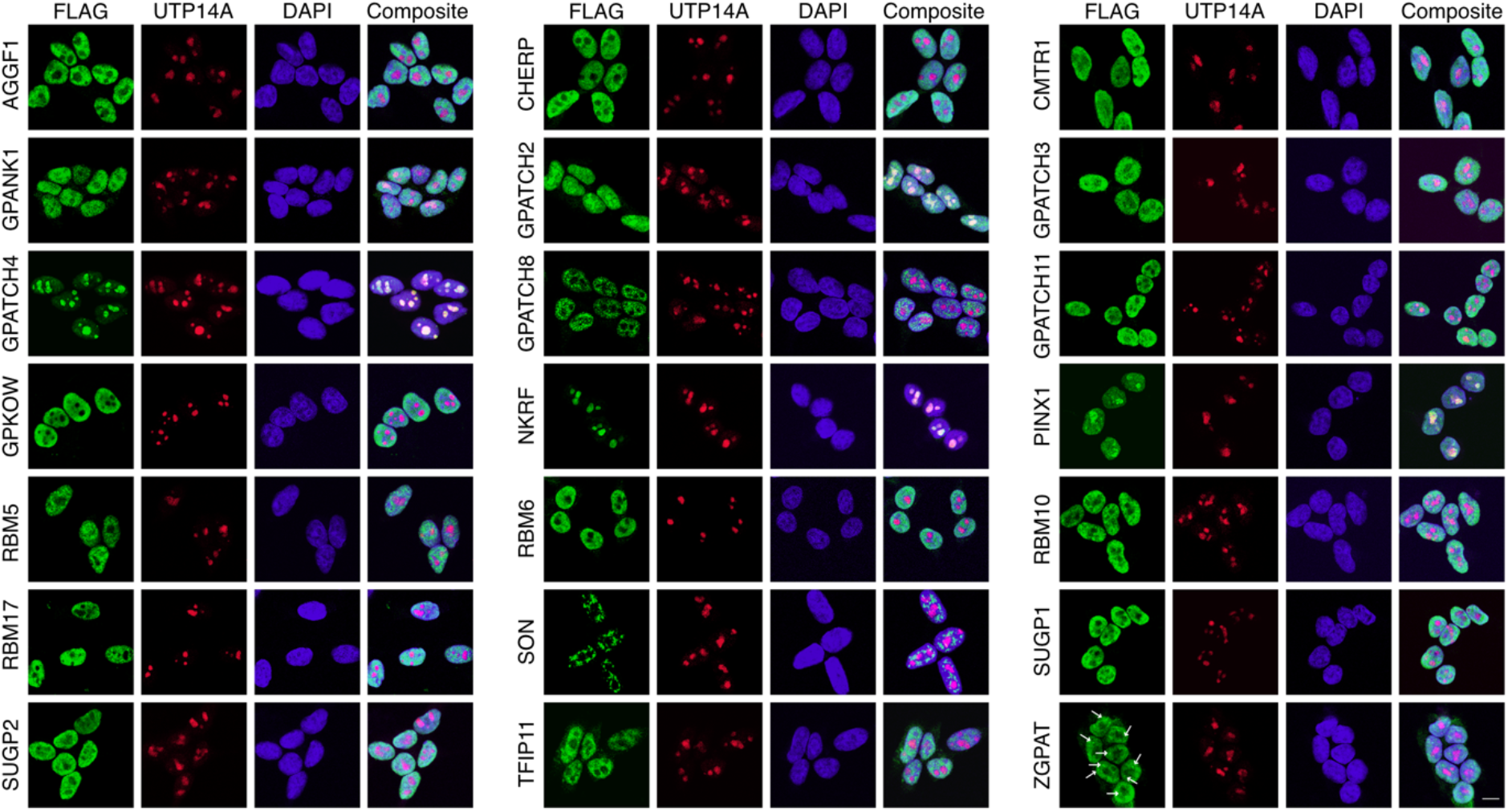
Sub-cellular localizations of human G-patch protein. FLAG-tagged G-patch proteins expressed from the stable cell lines were detected by immunofluorescence microscopy using an antibody against the FLAG tag (green). Nucleoli were detected using an antibody against UTP14A (red) and nuclear material was visualized by DAPI staining (blue). Overlays of the signals from the three channels are shown (Composite). All images are shown at the same scale and the scale bar represents 10 μm.

**Figure 4.**
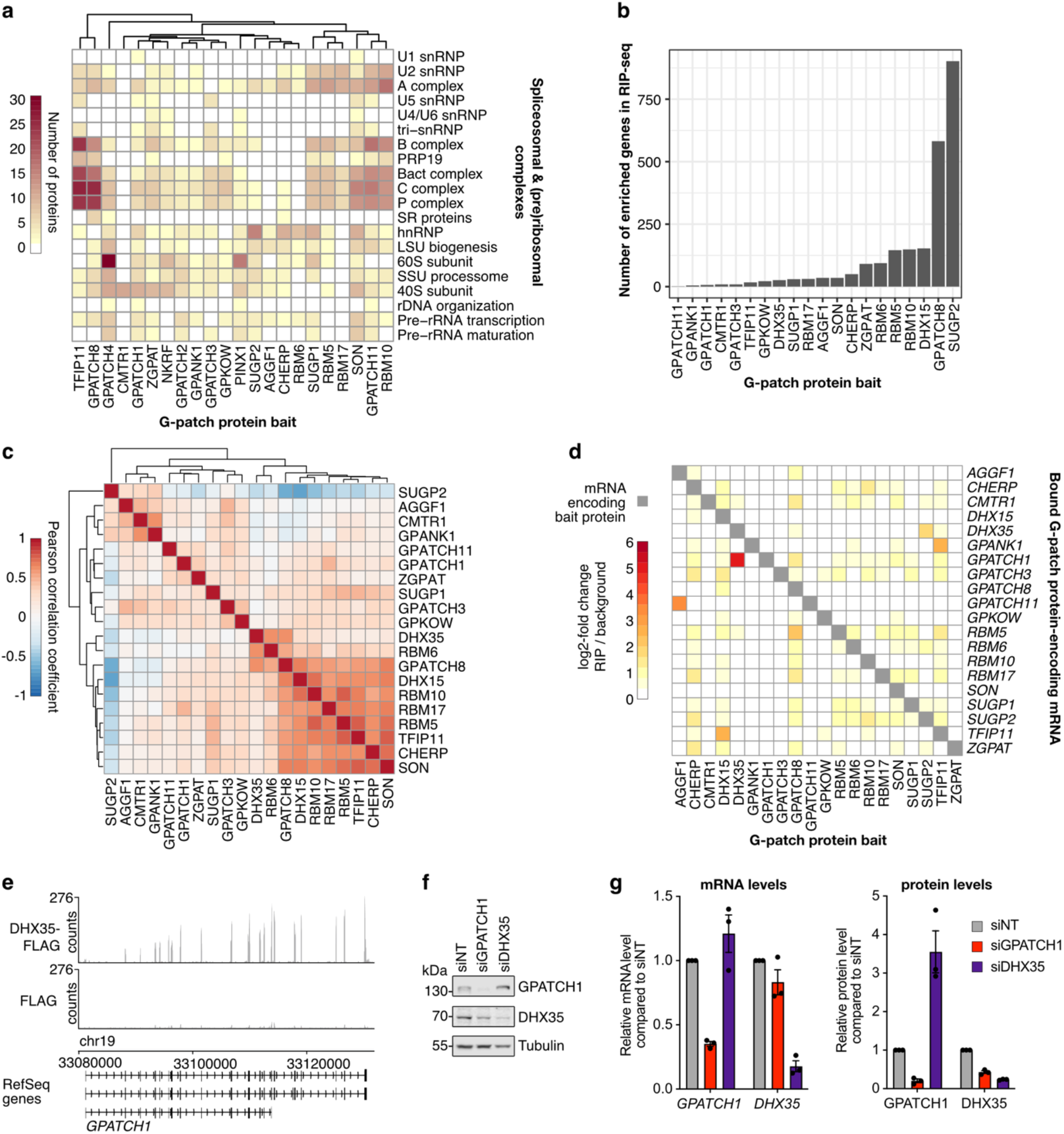
RNA interactomes of G-patch proteins and cross-regulation of DHX35 and GPATCH1 expression. **(a)** Heatmap representing the numbers of proteins in different spliceosomal complexes and (pre-)ribosomal complexes co-precipitated with each of the G-patch proteins by IP-MS. **(b)** RNA immunoprecipitation and sequencing (RIP-seq) of FLAG-tagged G-patch proteins/RNA helicases yields a range of bound RNAs with each the G-patch protein/RNA helicase compared to the FLAG control. **(c)** Heatmap showing the Pearson correlation coefficients of RIP-seq enrichments for genes enriched in at least one condition. **(d)** Heatmap showing the enrichment of *G-patch protein*/*DHX15*/*DHX35* mRNAs in the RIP-seq datasets. **(e)** Read coverage for DHX35-FLAG RIP-seq and the FLAG control over the *GPATCH1* gene. **(f-g)** Cells were transfected with non-target siRNAs (siNT) or those targeting GPATCH1 (siGPATCH1) or DHX35 (siDHX35). The levels of the indicated mRNAs were determined by RT-qPCR. The corresponding protein levels were analyzed by western blotting and quantified in three independent experiments. Data are presented as mean of n=3 experiments ± standard error.

The human G-patch proteins AGGF1, CHERP, CMTR1, GPANK1, GPATCH3, GPATCH11, GPKOW, RBM5, RBM6, RBM10, RBM17, SUGP1, SUGP2 and TFIP11 were all present in the nucleoplasm and excluded from nucleoli (**Fig. 3**). As previously observed, the well-characterized SR protein SON was concentrate in large nucleoplasmic foci representing splicing speckles^35,52^. Interestingly, GPATCH8 also displayed punctate signals distributed across the nucleoplasm, but these foci were much smaller than those of SON (**Fig. 3**). In addition to a diffuse nucleoplasmic signal, ZGPAT was also concentrated in small, nucleolar-adjacent nucleoplasmic foci likely representing Cajal bodies (**Fig. 3**), consistent with the detection of this protein in 35S tri-snRNP complexes^39^.

### RNA interactomes of human G-patch proteins reveal a broad range of RNA targets and regulation of GPATCH1 expression by DHX35

The nucleoplasmic localization of the majority of the human G-patch proteins suggests that they are involved in the early stages of gene expression. Further analysis of the protein interactomes of the G-patch proteins determined by IP-MS (**Supplementary Data 1**) highlighted the associations with numerous splicing factors (**Fig. 4a**). Strikingly, SR proteins were co-precipitated by GPATCH8, whereas hnRNP proteins were recovered with several G-patch proteins, especially SUGP2 (**Fig. 4a**). Robust associations of TFIP11, GPATCH8, SUGP1, RBM5, RBM10, SON, and GPATCH11 with the spliceosomal B, B^act^, C, and P complexes were observed. In addition, U2 snRNP and A-complex factors were enriched among proteins associated with a subset of these factors, including SUGP1, RBM5, RBM10, SON, and GPATCH11, as well as RBM17 and CHERP (**Fig. 4a**). In order to gain further insight into the pre-mRNA targets of the G-patch proteins, RNA immunoprecipitation coupled to deep sequencing (RIP-seq) was performed using extracts from the cell lines expressing FLAG-tagged G-patch proteins. NKRF, GPATCH4, GPATCH2 and PINX1 were excluded due their known/anticipated functions in ribosome assembly, and the RNA helicases, DHX15 and DHX35 were included for comparison.

Analysis of the RNAs enriched with each of these proteins revealed a broad range of numbers of transcripts recovered with each G-patch protein as well as differences in the types of RNA (**Fig. 4b**; **Supplementary Data 5**). GPATCH11, GPANK1, GPATCH1 and GPATCH3, for example, recovered only minimal numbers RNAs whereas >500 transcripts were co-purified with both SUGP2 and GPATCH8 (**Supplementary Fig. S2a**). While this may reflect variations in the nature of the associations of different G-patch proteins with cellular RNAs (direct vs indirect, stable vs transient, etc.), it also probably indicates that some G-patch proteins fulfil very specific roles, influencing the expression of a limited number of genes, whereas others likely have more pleiotropic functions, contributing to regulation of the expression of numerous RNAs. While GPATCH1 and DHX35 recovered comparable numbers of RNAs, indicative of their co-functionality, the amounts of RNAs bound by the DHX15-interacting G-patch proteins GPATCH8 and SUGP2 exceeded the number co-purified with the helicase (**Fig. 4b**), suggesting that these G-patch proteins may, at least partially, fulfil helicase-independent functions. Systematic correlation analysis of the RNA targets of each of the G-proteins and helicases highlighted a cluster of G-patch proteins (GPATCH8, RBM10, RBM5, RBM17, CHERP, TFIP11 and SON) that showed strong similarities to each other and to their cognate helicase DHX15 (**Fig. 4c**). In this analysis, SUGP2 was a clear outlier and strikingly, of the multiple RNAs recovered with SUGP2, 98% were unique to this G-patch protein (**Supplementary Fig. S2b**; see below for further details).

The identification of numerous RIP-seq targets raised the question whether there is cross-talk between any of the G-patch proteins and/or their cognate RNA helicases. Intriguingly, focused analysis of the mRNAs encoding the G-patch proteins/helicases revealed strong enrichment of the *GPATCH1* mRNA with the GPATCH1-associated RNA helicase DHX35 (**Fig. 4d, e**). To investigate the consequences of DHX35 association with the *GPATCH1* mRNA, RNAi-mediated depletion of DHX35, and for comparison GPATCH1, was established (**Fig. 4f**). On the RNA level, depletion of DHX35 mildly, but not significantly, increased the level of the *GPATCH1* mRNA (**Fig. 4g, left**). However, on the protein level, depletion of DHX35 led to a >3-fold increase in GPATCH1 (**Fig. 4g, right**). Furthermore, despite the absence of GPATCH1–*DHX35* mRNA interactions (**Fig. 4d**) and no significant decrease in the level of the *DHX35* mRNA in cells lacking GPATCH1, depletion of GPATCH1 lead to a >2-fold decrease in DHX35 protein (**Fig. 4g, right**). These data suggest that i) binding of DHX35 to the *GPATCH1* mRNA regulates expression of the GPATCH1 protein and ii) formation of a complex with GPATCH1 stabilizes the RNA helicase DHX35.

### G-patch proteins associate with diverse classes of RNAs and ZGPAT is required for efficient 2′-*O*-methylation of the U2 and U5 snRNAs

Next, the RIP-seq datasets were evaluated considering the classes of RNAs associated with each G-patch protein. While a substantial fraction of the transcripts retrieved with several G-patch proteins, including GPKOW, AGGF1, RBM6, SUGP1 and SUGP2, were long non-coding RNAs (lncRNAs), most G-patch proteins predominantly associated with (pre-)mRNAs (**Fig. 5a**; **Supplementary Data 5**). Consistent with roles in pre-mRNA splicing, a subset of G-patch proteins also recovered snRNAs (**Fig. 5a**); alongside ZGPAT, this group includes CHERP and RBM17, which are known to associate with the U2 snRNP and the spliceosomal A complex (**Fig. 4a**)^44^, as well as TFIP11 that contributes to optimizing U4/U6.U5 snRNP assembly^53^.

**Figure 5.**
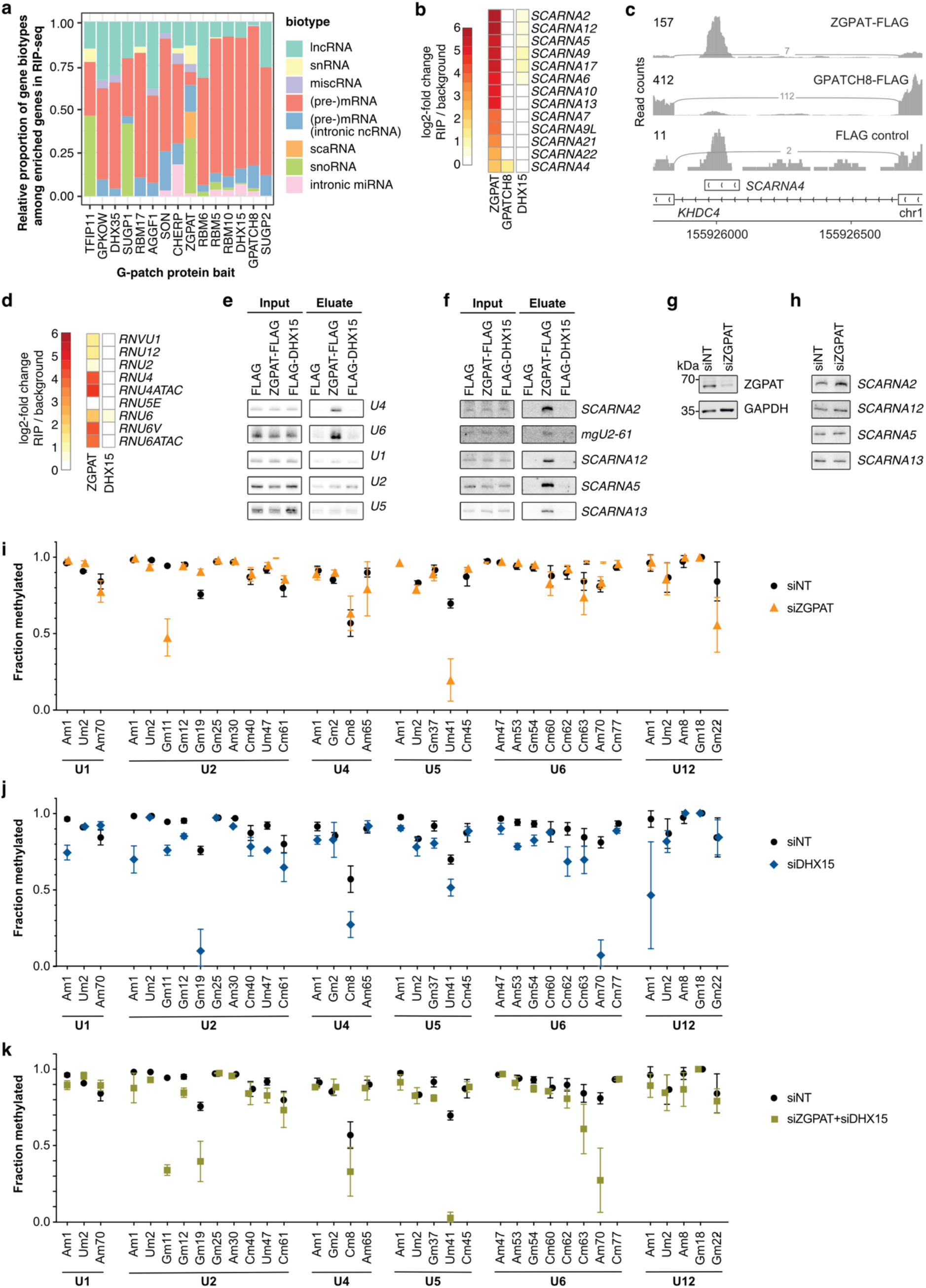
ZGPAT binds snRNAs and scaRNAs, and is required for efficient 2′-*O*-methylation of snRNAs. **(a)** Biotype distribution of RNAs >10 enriched with FLAG-tagged G-patch proteins/RNA helicases compared to the FLAG control as determined by RIP-seq. **(b)** Enrichment of different scaRNAs in the ZGPAT, GPATCH8 and DHX15 RIP-seq datasets are shown. **(c)** ZGPAT-FLAG, GPATCH8-FLAG and FLAG control RIP-seq coverage of the region of the KHDC4 gene surrounding SCARNA4. **(d)** Enrichment of different snRNAs in the ZGPAT and DHX15 RIP-seq datasets are shown. **(e)** ZGPAT or DHX15 RIP analyzed by northern blotting validates snRNA binding of ZGPAT. 1% of the input was loaded. **(f)** ZGPAT or DHX15 RIP analyzed by northern blotting validates scaRNA binding of ZGPAT. 1% of the input was loaded. **(g)** Western blot detecting ZGPAT and tubulin (loading control) levels upon cell treatment with non-target siRNAs (siNT) or those targeting ZGPAT. **(h)** ScaRNA levels detected by northern blot upon knock-down of ZGPAT or cell treatment with siNT. **(i-k)** RNAs from cells treated with siNT, siZGPAT (i), siDHX15 (j) or siZGPAT and siDHX15 (k) was used for RiboMeth-seq and the methylation scores for each 2′-*O*-methylated nucleotide in the U1, U2, U4, U5, U6 and U12 snRNAs is plotted. n=3 experiments were performed and data are presented as mean ± standard deviation.

Notably, reads mapping to genes encoding various other small non-coding RNAs (small nucleolar RNAs (snoRNAs), small Cajal body-associated RNAs (scaRNAs) and micro RNAs (miRNAs)) were also recovered with specific G-patch proteins (**Fig. 5a**; **Supplementary Data 5**). For example, RIP-seq for ZGPAT, TFIP11, SUGP1, and various other G-patch proteins (RBM5, RBM6, RBM10 and GPATCH8) recovered reads mapping to snoRNA genes (**Supplementary Fig. S3a**), and ZGPAT also enriched scaRNAs, consistent with its presence in Cajal bodies (**Fig. 3**; **Fig. 5a, b**). In humans, most sno/scaRNAs (>80-90%) and miRNAs (approx. 50-70%) are present within the introns of protein-coding genes^54,55^, raising the question whether these G-patch proteins bind to the excised ncRNAs or rather to sno/scaRNA/miRNA sequences within pre-mRNA introns. For the G-patch proteins recovering few snoRNAs/miRNAs (e.g. GPATCH8) the pre-mRNAs within which they are encoded were also enriched to a similar extent (**Supplementary Fig. S3b**), suggesting that the intron-containing pre-mRNA, rather than the ncRNA, is bound by the G-patch protein. This is in contrast to the G-patch proteins TFIP11 and ZGPAT where only a minor fraction of pre-mRNA transcripts containing the enriched ncRNAs were recovered, and the enrichment of these pre-mRNAs was much lower than the enrichment of the sno/scRNAs (**Supplementary Fig. S3b**), indicating specific association of the G-patch protein with the mature sno/scaRNA. This is well exemplified by the profiles of ZGPAT and GPATCH8 RIP-seq data on the *KHDC4* pre-mRNA, which contains SCARNA4 within intron 6; while the majority of reads derived from GPATCH8-associated RNAs mapped to the upstream/downstream exons and coverage across the intron was relatively uniform, for ZGPAT a well-defined peak on the intronic regions encoding the scaRNA was observed (**Fig. 5c**).

Alongside TFIP11 and ZGPAT, another G-patch protein for which a substantial portion of RIP-seq reads mapped to snoRNA genes was SUGP1 (**Fig. 5a**; **Supplementary Fig. S3a**). Strikingly, several of these snoRNAs (SNORD22, SNORD25, SNORD26, SNORD27, SNORD28 and SNORD30) are encoded within the snoRNA host gene *SNHG1* (**Supplementary Fig. S3a**), suggesting that SUGP1 plays a role in splicing of this lncRNA. This notion is supported by i) the recovery of sequences derived from these snoRNAs with other characterized splicing factors (RBM5, RBM6 and RBM10), and ii) the mapping profile of the SUGP1 RIP-seq data across *SNHG1*, which shows coverage across the transcript (**Supplementary Fig. S3a, c**). Among the snoRNAs encoded within *SNHG1* is SNORD27, which fulfils a non-canonical role in alternative splicing^56^. The strong enrichment of SNORD27 sequences with SUGP1 (**Supplementary Fig. S3a, c**) raises the possibility that SUGP1 may also associate with the mature form of the snoRNA in the context of SNORD27-mediated alternative splicing regulation, but this remains to be confirmed.

As ZGPAT showed the most diverse spectrum of associated RNA types, the functional relevance of these interactions was explored next. The RIP-seq datasets highlighted robust interaction of ZGPAT with the U4 and U6 snRNAs, but not U5 and this was confirmed by northern blotting (**Fig. 5d, e**). These interactions are consistent with the association of ZGPAT with a ∼35S U4/U6 snRNP complex into which to U5 snRNP is assembled^39^. Analysis of the scaRNAs enriched with ZGPAT highlighted SCARNA2, SCARNA12, SCARNA5, SCARNA9, SCARNA17 and SCARNA6 as >5-fold enriched with ZGPAT compared to the control (**Fig. 5b, f**). In line with the absence of scaRNA enrichment with DHX15 observed by RIP-seq, northern blotting also confirmed the association of ZGPAT with scaRNAs to be specific to the G-patch protein (**Fig. 5b, f**). The impact of RNAi-mediated ZGPAT depletion (**Fig. 5g**) on scaRNA levels was analyzed, and no significant differences were observed (**Fig. 5h**). As scaRNAs guide snRNA modifications^57^, the RiboMeth-seq approach, which enables simultaneous detection of all 2′-*O*-methylations (Nm) in abundant RNA species^58,59^, was applied to examine how lack of ZGPAT and/or DHX15^16,17^ affect snRNA 2′-*O*-methylation (**Supplementary Data 6)**. Compared to control cells, in those lacking ZGPAT, U2-Gm11, U5-Um41 and U12-Gm22 were markedly reduced whereas methylation of the normally sub-stoichiometric U2-Gm19 was elevated (**Fig. 5i**). Strikingly, the scaRNAs guiding these methylations (U2-Gm11 – SCARNA2, U2-Gm19 – SCARNA9, U5-Um41 – SCARNA5/SCARNA6, U12-Gm22 – SCARNA17) were among those most enriched with ZGPAT in the RIP experiments (**Fig. 5b**). Interestingly, U2-Gm11 and U5-Um41 were also reduced in cells lacking DHX15 and in contrast to cells depleted of ZGPAT, U2-Gm19 was decreased (**Fig. 5j**). Depletion of DHX15 also impaired efficient methylation of several other sites across the snRNAs, but these effects were independent of ZGPAT. Combined lack of ZGPAT and DHX15 exacerbated the methylation defects at sites U2-Gm11 and U5-Um41 (**Fig. 5k**), potentially implying that, rather than functioning cooperatively, the G-patch protein and RNA helicase may be independently required to ensure efficient 2′-*O*-methylation of these nucleotides. This notion is supported by the opposing effects of depletion of ZGPAT and DHX15 on U2-Gm19 (**Fig. 5i-k**).

### DHX15 and many of its G-patch protein cofactors concertedly regulate alternative pre-mRNA splicing and mRNA expression

The association of numerous G-patch proteins with splicing factors (**Fig. 4a**) and pre-mRNAs (**Fig. 5a**) is in line with roles in mRNA maturation so RNA-seq was applied to compare the alternative splicing and gene expression landscapes of cells lacking individual G-patch proteins (**Supplementary Data 7**). RNAi-mediated depletion of the nucleoplasmic G-patch proteins and DHX15 was established, and reduction of target mRNA levels was confirmed by RT-qPCR and in the respective total RNA-seq datasets (**Fig. 6a**; **Supplementary Fig. S4a**). Alternative splicing (AS) patterns (**Supplementary Fig. S4b**) were then investigated using rMATS for skipped exons (SE), alternative 3′ or 5′ splice site usage (A3SS and A5SS) and mutually exclusive exons (MXE). To quantify intron retention across annotated introns of diverse sizes, intron retention (IR) was measured analogously to the approach taken for calculating splicing efficiencies in FRASER or SPI^60–62^. To allow for comparison in the degree of alternative splicing changes by G-patch/DHX15 knock-down, only events that passed all applied cutoffs in 80% of the datasets were considered for further analysis. Quantified splicing events originated from approximately 12,000 genes, of which 4-19% of genes contained significant AS events (**Fig. 6b**). The knock-down of SON and DHX15 yielded, with more than 3,000 events, most significant AS events (**Fig. 6b-c**). Depletion of AGGF1, GPATCH8, GPANK1 or ZGPAT yielded less exon inclusion overall, whereas reduction of SON, DHX15, SUGP1, RBM5, RBM10 and RBM17 led to the reverse trend (**Fig. 6b, d**). IR forms the largest group of significant AS events (**Fig. 6c, Supplementary Fig. S4c**), but the fraction of significant AS events relative to all quantified events is comparable to other types of AS (see dot sizes in **Fig. 6b**). Most knock-downs yield increased IR, except for SUGP2, CMTR1 and ZGPAT (**Fig. 6b**).

**Figure 6.**
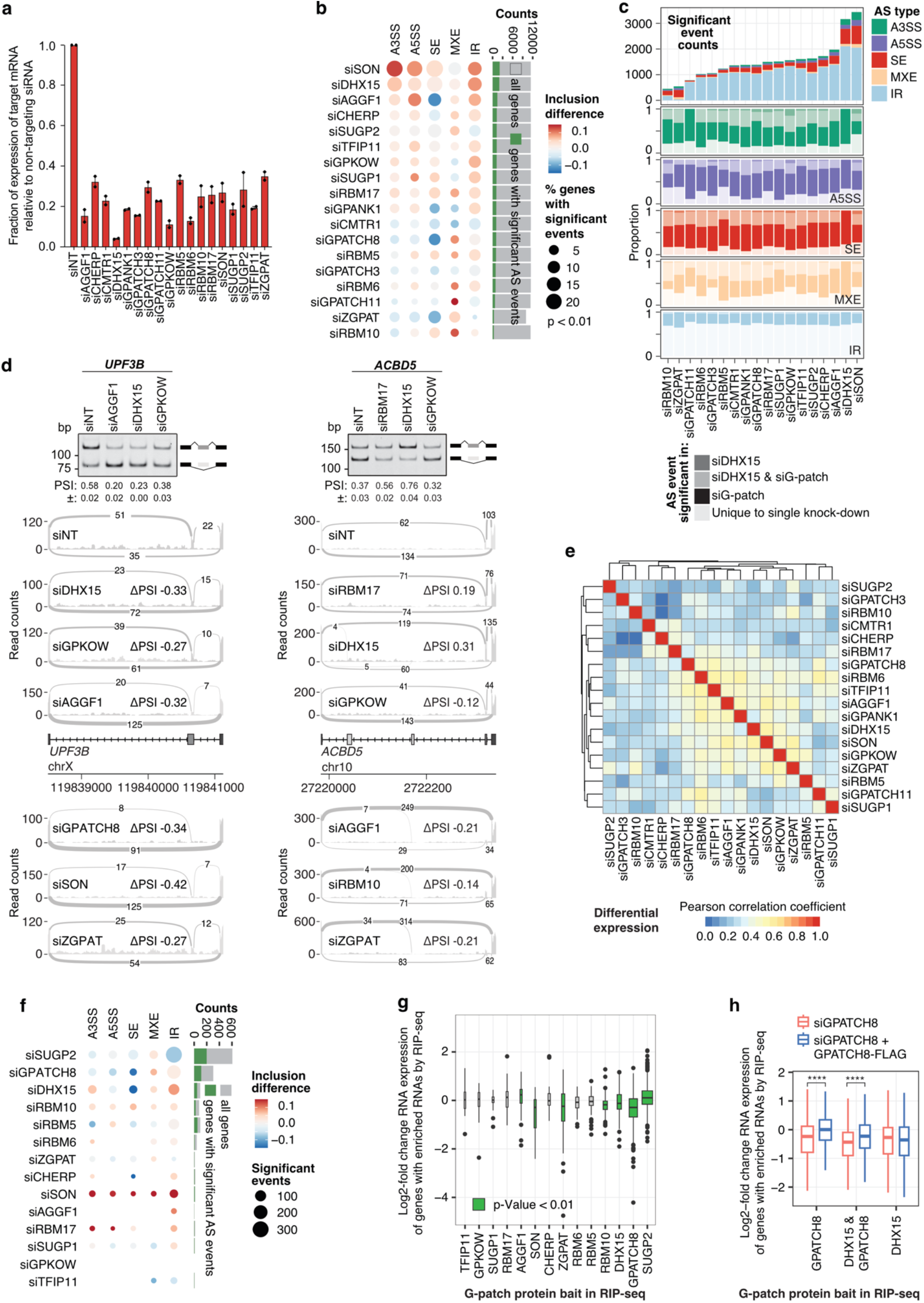
Overlapping effects on alternative splicing and gene expression by multiple G-patch proteins. **(a)** Cells were transfected with non-target siRNAs (siNT) or those targeting the indicated G-patch proteins, and the levels of the target mRNAs, relative to siNT, after normalization to *EMC7*, were determined by RT-qPCR in n=2 biological replicates. **(b)** Different extent and directionality of global alternative splicing changes for groups of G-patch proteins/ DHX15. Dotplot of median inclusion difference for significant AS events (p-value < 0.01) and barplot of gene counts with and without significant AS events. Each RNA-seq dataset consists of n=3 biological replicates. **(c)** Absolute counts of significant AS events per knock-down and proportion of overlap in events with other G-patch and *DHX15* mRNA knock-downs. **(d)** RT-PCR validation of two significant SE events and corresponding sashimi plots for probed knock-downs (above gene diagram) and knock-downs that were not assayed by RT-PCR (below gene diagram). Sashimi plots reflect the cumulative coverage of n=3 biological replicates per condition. siNT coverage reflects a representative pool of three siNT replicates out of nine. For RT-PCR experiments, n=2 experiments were performed and percent spliced in (PSI) is given ± standard error of mean. **(e)** Heatmap illustrating degree of correlation of differential gene expression, relative to the non-target siRNA control, between siRNA knock-downs. **(f)** As (b), but for genes encoding RNAs enriched by RIP-seq. **(g)** Boxplot showing the distribution of differential gene expression relative to the non-target siRNA control for genes encoding RNAs enriched by RIP-seq. If differential expression relative to genes encoding RNAs not enriched by RIP-seq in each knock-down is significantly up or down, boxes are colored green (two-sided Wilcoxon rank sum test with Benjamini & Hochberg correction, p-values: AGGF1 0.008, DHX15 0.006, RBM5 4.7e-06, RBM10 3.3e-14, SON 0.001, SUGP2 1.6e-25, ZGPAT 0.004). **(h)** Rescue of differential expression in *GPATCH8* knock-down of GPATCH-bound and/or DHX15-bound RNAs with expression of siRNA-insensitive, FLAG-tagged GPATCH8 protein. Two-sided Wilcoxon rank sum test with Benjamini & Hochberg correction, **** p-values < 0.0001, GPATCH8 2.7e-20, GPATCH8 & DHX15 5.5e-11.

Given that DHX15 interacts with most G-patch proteins (**Fig. 1c-e**), a substantial overlap or co-regulation of alternative splicing events by DHX15 and G-patch proteins was expected. Indeed, 76% of significant non-IR DHX15 events were also differentially spliced upon knock-down of one or more G-patch proteins (**Fig. 6c**). Also, the overlap of significant non-IR events of individual G-patch knock-downs to others was in a similar range, suggesting extensive crosstalk and co-regulation of alternative splicing (**Fig. 6c**). Several events that showed differential SE in multiple knock-down conditions were corroborated using RT-PCR (**Fig. 6d, Supplementary Fig. S4d**). To investigate how individual alternative splicing patterns compared across G-patch protein depletion, Pearson correlations of ΔPSI (percent spliced in) values across all non-IR events that were quantified as significantly different in at least one condition were performed (**Supplementary Fig. S4e**). Depletion of G-patch proteins, including TFIP11, GPATCH8, GPATCH11, SUGP1, that grouped together by their degree of binding to spliceosomal subcomplexes (**Fig. 4a**) also exhibited more similar splicing profiles. Of all knock-downs, GPATCH8 depletion showed the highest correlation coefficients to other knock-downs (**Supplementary Fig. S4f**), which is reminiscent of its RNA binding profile that displayed high similarity to other G-patch protein interactomes in RIP-seq (**Supplementary Fig. S2b**). IR, however, showed a small overlap to other G-patch or DHX15 knock-downs, suggesting that general splicing efficiency changes are more specific to individual perturbations of G-patch protein or DHX15 levels (**Fig. 6c**).

The observed changes in alterative splicing may affect mRNA levels so the correlation of differential expression profiles upon G-patch protein or DHX15 depletion was investigated (**Fig. 6e**). Similar to our observation by RIP-seq (**Fig. 4c, Supplemental Figure S2b**), the differential expression observed in cells lacking SUGP2 largely differed to most other knock-downs. Also analogous to the RNA interactomes, differential expression in several G-patch protein depletions, including GPATCH8, correlated equally well and showed similarity to DHX15.

SUGP2, GPATCH8 and DHX15 recovered the highest amount of RNA targets in RIP-seq (**Fig. 4b**), so focusing on AS and differential expression changes of bound RNAs can provide insights into the direct roles of these proteins in gene expression. For SUGP2-bound RNAs, depletion of SUGP2 decreased intron retention, suggesting a repressive role of SUGP2 on pre-mRNA splicing (**Fig. 6f**). RNAs bound by GPATCH8 and DHX15 showed a common trend of increased exon skipping upon depletion of either of the G-patch protein or the RNA helicase (**Fig. 6f**). Similar to the effects of G-patch protein depletion on overall AS (**Fig. 6b**), when considering AS in the sub-population of RNAs bound by each G-patch protein, only few AS events changed significantly upon G-patch protein depletion. This suggests that binding of G-patch proteins may also influence other aspects of mRNA expression. Nevertheless, depletion of GPATCH8, DHX15, RBM10, ZGPAT or SON significantly decreased the levels of transcripts bound by the corresponding G-patch proteins, and mRNAs bound by SUGP2 or AGGF1 showed increased expression upon depletion of the G-patch protein. Complementation experiments in which siRNA-insensitive, FLAG-tagged GPATCH8 was expressed in cells depleted of the endogenous protein (**Supplementary Fig. S4g**) showed that the mRNA expression differences observed in GPATCH8-bound RNAs upon depletion of the G-patch protein were rescued by re-expression of the FLAG-tagged protein (**Fig. 6h**). This was also the case for RNAs bound by both GPATCH8 and DHX15, but not for RNAs associated with only DHX15 (or DHX15 and other G-patch proteins), implying that the observed differences in gene expression arise from a direct effect of GPATCH8–DHX15 on their bound RNAs. Taken together, the family of G-patch proteins and DHX15 influence thousands of alternative splicing events and regulate RNA expression, with common genes, introns and exons often targeted, supporting the notion of functionality as an interconnected network.

### DHX15-independent suppression of pre-mRNA splicing by SUGP2 intron binding

Throughout our analyses, SUGP2 has emerged as outlier G-patch protein that affects RNA expression distinctly from other G-patch proteins, and mostly independent of DHX15. The immunoprecipitation experiments identified little DHX15-bound SUGP2 protein (**Fig. 1c-e**) and its G-patch only very weakly stimulated the ATPase activity of DHX15 (**Fig. 2e**), without promoting RNA binding by DHX15 (**Fig. 2k**). Furthermore, its association with 12 hnRNP proteins and only a few A complex proteins is unique to this G-patch protein (**Fig. 4a**). Also, SUGP2 RIP-seq showed a very different enrichment profile compared to other G-patch proteins (**Fig. 4c, Supplementary Fig. S2b**) and in addition to (pre-)mRNAs, several lncRNAs were robustly recovered together with SUGP2 (**Fig. 5a**). Non-IR alternative splicing changes follow the direction of changes in the DHX15 knockdown, but are typically much weaker (**Fig. 6b, Supplementary Fig. S4c-d**), and we speculate that these effects may be influenced by the small population of SUG2–DHX15 complexes. Interestingly, IR is relieved rather than increased upon SUGP2 knock-down (**Fig. 6b, f, Supplementary Fig. S4c**) and global gene expression changes upon SUGP2 knock-down were distinct from other G-patch protein or DHX15 knock-downs (**Fig. 6e, g**), raising the possibility of a DHX15-independent role for SUGP2 in suppressing splicing. This notion is consistent with the association of SUGP2 with hnRNP proteins, which are broadly implicated in making splice sites more or less accessible to the spliceosome^63^.

Motivated by these results, further analyses were performed to probe the features of SUGP2-bound RNAs and clarify the role of SUGP2 in gene expression. Combining our RIP-seq data with published eCLIP data^64^, a strong overlap in bound RNAs and enrichment of intronic signal was observed (**Fig. 7a,b**). This is consistent with the characterization of SUGP2 as deep intronic RNA binding protein^64,65^. Intronic signal was highest for SUGP2-bound RNAs identified by RIP-seq (**Fig. 7b**). The FLAG control and total RNA-seq data for these genes also showed high intronic signal compared to an equally sized, randomly chosen gene set (**Fig. 7b**), indicating overall lower splicing efficiency of SUGP2-bound introns than those not associated with SUGP2. As observed by global AS analysis (**Fig. 6b**), splicing efficiency increases upon knock-down of SUGP2, such that IR modestly decreases (**Fig. 7c**). This effect was rescued by the expression of full-length FLAG-tagged SUGP2 protein (**Fig. 7d; Supplementary Fig. S5a**).

**Figure 7.**
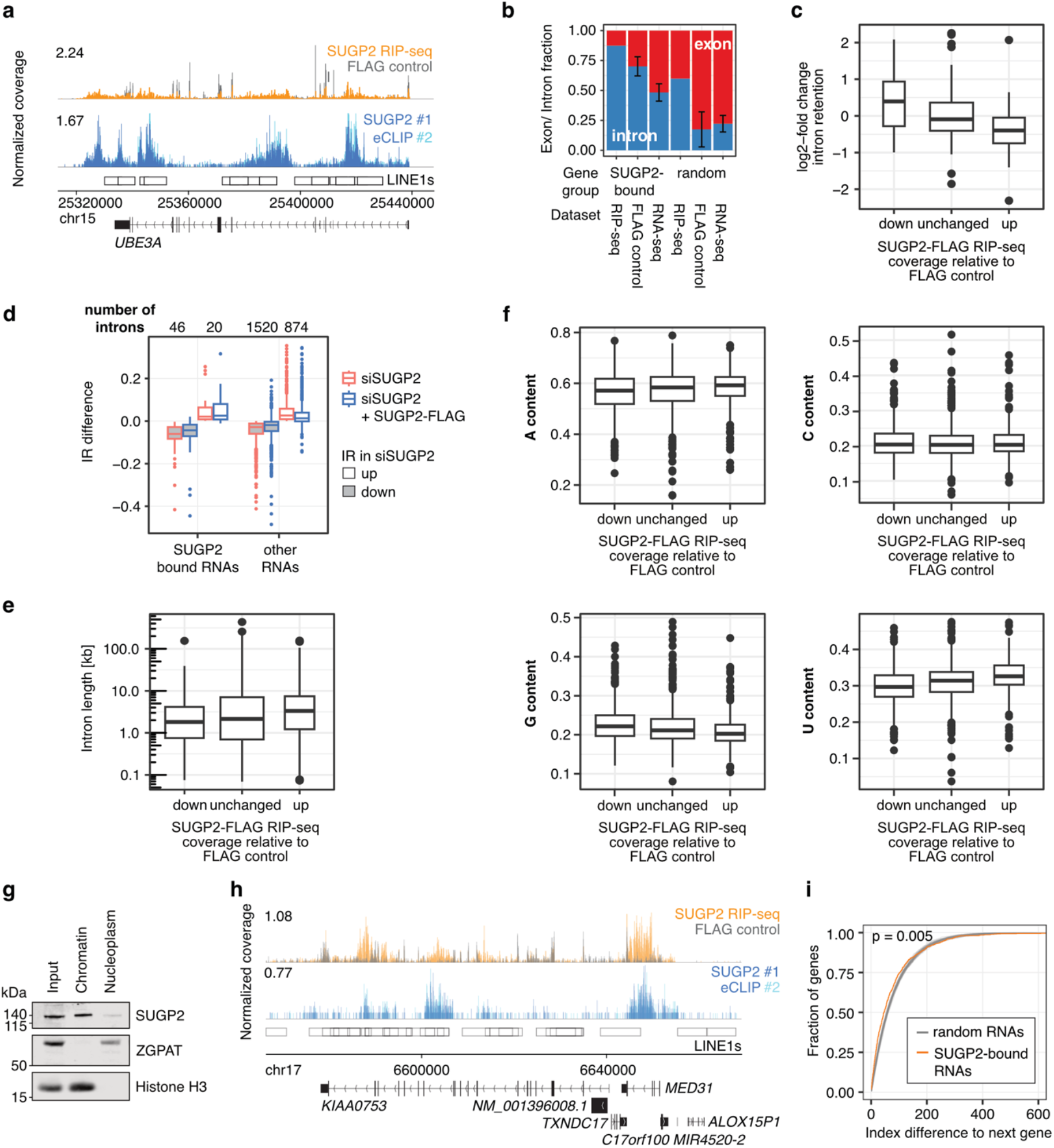
Chromatin-associated SUGP2 reduces splicing efficiency and binds to long U-rich introns in transcripts of often adjacent genes. **(a)** Read coverage for SUGP2-FLAG RIP-seq, FLAG control and SUGP2 eCLIP with control over the *UBE3A* gene. LINE1s reflect collapsed LINE1 element annotations. **(b)** Fraction of reads mapping to exons and introns for SUGP2 bound RNAs or randomly chosen expressed RNAs for different datasets. Error bars reflect SD after bootstrapping (100 iterations). **(c)** Intron retention differences in groups defined in (c). Kolmogorov-Smirnov test with Benjamini & Hochberg correction, p-Value 0.026 between ‘unchanged’ and ‘up’). **(d)** Rescue of intron retention pattern in *SUGP2* knock-down of SUGP2-bound and other RNAs with expression of siRNA-insensitive, FLAG-tagged SUGP2 protein. Two-sided Wilcoxon rank sum test with Benjamini & Hochberg correction, p-Value: other RNAs / down 5.85e-22, other RNAs / up 1.68e-24. **(e)** Distribution of intron lengths grouped by signal in SUGP2 RIP-seq of the respective introns. Two-sided Wilcoxon rank sum test with Benjamini & Hochberg correction, p-Value: unchanged / up 0.009, unchanged / down 0.017, up / down 6.07e-08. **(f)** Nucleotide composition of introns based on the grouping in (e). Two-sided Wilcoxon rank sum test with Benjamini & Hochberg correction, p-Value: A unchanged / down 0.007, A up / down 0.0001, G unchanged / up 3.0e-05, G unchanged / down 3.6e-05, G up / down 1.7e-13, U unchanged / up 1.5e-08, U unchanged / down 1.5e-08, U up / down 1.0e-22. **(g)** Nuclei extracted from HEK293 cells were separated into nucleoplasm and chromatin fractions, and chromatin-bound proteins were released by DNase treatment. Proteins from the total cell extract (input), and nucleoplasm and chromatin fractions were analysed by western blotting using the indicated antibodies. Representative data from n=3 experiments are shown. **(h)** Read coverage for SUGP2-FLAG RIP-seq, FLAG control and SUGP2 eCLIP with control for adjacent genes. LINE1s reflect collapsed LINE1 element annotations by RepeatMaskerViz v4.0.7. **(i)** Cumulative distribution of gene index differences for SUGP2 bound RNAs relative to randomly selected RNAs and their respective gene index differences (50 iterations displayed; two-sided Wilcoxon rank sum test with Benjamini & Hochberg correction, adjusted p-value reflects mean of comparison to random controls). Gene index reflects rank by the mid-gene position across each chromosome for same stranded non-overlapping genes.

SUGP2-bound introns are distinct in their architecture; they do not show differences in their canonical splice site strength (**Supplementary Fig. S5b**) but are longer than non-enriched introns, and are depleted of guanines and enriched in uridines (**Fig. 7e, f**). The robust binding of SUGP2 to intronic regions, raises the possibility that RNA binding and splicing regulation could occur, or be initiated, co-transcriptionally. Indeed, cell fractionation experiments identified the majority of SUGP2 in the chromatin-associated fraction (**Fig. 7g**). Moreover, SUGP2 RIP-seq and eCLIP data suggest more association to transcripts of adjacent genes than expected by chance (**Fig. 7h, i**), consistent with the notion of nascent RNA binding within the context of particular genomic regions. Altogether, these results indicate that SUGP2 binding suppresses intron excision, potentially by sterically impeding access of splicing factors, a mode of RNA expression modulation distinct from other G-patch proteins.

## DISCUSSION

The G-patch motif, first recognized in the splicing factor SON and subsequently found to be present in > 20 human proteins, is emerging as a distinctive feature of proteins involved in various different aspects of gene expression^9,10,41^. Consistent with this, over the course of evolution from yeast to humans, the number of G-patch proteins has increased approximately four-fold, likely reflecting the increased fine-tuning and regulation of gene expression in higher eukaryotes. The identification of cognate RNA helicases for each human G-patch protein highlighted specific G-patch protein–RNA helicase partnerships (GPKOW–DHX16 and GPATCH1–DHX35) as well as an extensive network of G-patch proteins centred around the multifunctional RNA helicase DHX15. By systematically comparing the effects on RNA helicase activity, sub-cellular localizations, protein and RNA interactomes, and influences on the transcriptome across the network of human G-patch proteins, this study highlights diverse ways in which G-patch proteins contribute to mRNA maturation.

Cofactor proteins can direct helicases to their RNA targets as well as promote or supress helicase activity either allosterically or by modulating their affinities for substrates^25,66^. The G-patch proteins are ideally constructed to coordinate recruitment of their cognate helicases to appropriate substrates with helicase activation by combining protein–protein/–nucleic acid interaction domains with the G-patch motif (**Fig. 1a**). In the experimentally determined NKRF G-patch–DHX15 structure^20^, G-patch binding tethers the two RecA-like domains of the helicase to constrict the RNA-binding channel^20^. Single-molecule FRET experiments with Prp43 and Pfa1 (yeast homologues of DHX15 and GPATCH2) have also demonstrated that G-patch binding opens the catalytic cleft to facilitate ADP-ATP exchange upon ATP hydrolysis^18^. Our *in vitro* analyses reveal that most DHX15-associating G-patches stimulate RNA binding and ATP hydrolysis by the helicase (**Fig. 2e, k**). Hence, the principles of helicase activation by cofactor-driven enhanced RNA binding stimulating ATPase activity and helicase processivity that were observed for individual examples previously apply broadly across the G-patch protein family. Nevertheless, comparing DHX15 activation by each G-patch under identical conditions demonstrated a broad range of extents of ATPase stimulation (from ∼1.5–7-fold; **Fig. 2e**), despite high conservation of the G-patch motifs on the amino acid sequence level (**Supplementary Fig. S1a**). In cells, it is likely that regions of the cofactors outside the G-patch also contact DHX15, potentially enhancing the stimulatory effects of the G-patches alone^67,68^. However, it is also possible that the variable degrees of DHX15 stimulation by the different G-patch proteins reflect differences in the extent of helicase activity required on the substrates of each G-patch protein–RNA helicase complex. In the yeast G-patch proteins Pxr1 and Tma23, the activating G-patch has been shown to co-exist alongside an inhibitory element termed the I-patch that restrains Prp43 ATPase activity^69^. While such elements are yet to identified in human G-patch proteins, this raises the possibility that the disparate degrees of DHX15 activation observed for the different human G-patch proteins are balanced by other regulatory elements within the G-patch proteins. Strikingly, no stimulation of DHX15 was observed by the G-patch of RBM6. The mechanistic basis for this remains unclear, but it is possible that RBM6 functions cooperatively with the highly related and closely associated RBM5^42^, which strongly stimulates DHX15 ATPase activity (**Fig. 2e**). Alternatively, RBM6 could potentially act as a placeholder, blocking the interaction of DHX15 with other G-patch cofactors, thus sequestering the helicase in a non-productive state.

The RNA helicase interactions (**Fig. 1e**) and stimulatory effects of the G-patches on RNA helicase ATPase activity (**Fig. 2e**) are highly specific, raising the question how this specificity is achieved. Mutational analysis of DHX35 and GPATCH1 suggest a mode of helicase– cofactor interaction analogous to the experimentally determined RNA helicase–G-patch structures^19,20^, implying that these RNA helicase–G-patch protein complexes share a common interaction interface. However, in contrast to DHX15, the ATPase activities of neither DHX16 nor DHX35 were strongly stimulated by their associated G-patch proteins *in vitro* (**Fig. 2f-i**). In yeast, association of the DHX16 homologue Prp2 with spliceosomes promotes ATP hydrolysis and the presence of Spp2 (yeast homolog of GPKOW) within these complexes then converts this catalysis into a structural remodeling activity^70^. It is possible that similar principles also rationalize the lack of ATPase activity observed for the purified DHX35–GPATCH1_GP_ complex on a model RNA substrate. Strikingly, the non-DHX15-associated G-patches of GPKOW and GPATCH1 are the only G-patches with the ability to bind RNA (**Fig. 2j**). These distinct properties of the GPKOW–DHX16 and GPATCH1–DHX35 complexes, compared to DHX15 and its network of G-patch protein cofactors, may reflect adaptations of the core “DHX15–G-patch” mode of action that optimize these unique complexes for their specific cellular functions. In the case of GPKOW, *in vitro* RNA binding ability is in line with structural evidence showing that its yeast counterpart Spp2 tethers its cognate helicase to pre-mRNA substrates within B^act^ complexes to facilitate 3′-5′ translocation of the helicase^71^.

In *C. thermophilum*, the Dhx35–Gpatch1 complex has recently been implicated in the rejection of aberrant splicing substrates prior to the first splicing reaction^47^. Interestingly, Gpatch1 recruits Dhx35 to the U2 snRNA within a complex where Dhx15 is bound to the 3′ end of the U6 snRNA, demonstrating that different RNA helicase–G-patch protein complexes can function synergistically on a common RNP substrate. As human DHX35 and GPATCH1 have been identified as components of the spliceosomal complex C^72,73^, it is likely that their function in spliceosomal quality control is conserved. Our data reveal that the co-functionality of GPATCH1 and DHX35 is underpinned by a regulatory circuit, spanning the RNA and protein levels, in which the helicase and cofactor influence the expression of each other. DHX35 robustly binds the *GPATCH1* mRNA and suppresses its expression (**Fig. 4d-g**), likely to ensure that both proteins are present in cells at appropriate levels. This notion is further supported by the finding that formation of a complex with GPATCH1 stabilizes DHX35. Among the G-patch family, RNA helicase-mediated regulation of G-patch protein expression seems to be unique for DHX35–GPATCH1; such a mode of interdependent regulation would not be appropriate for DHX15 due to its multifunctionality and interactions with many cofactor proteins that each require to be expressed at a particular level.

Although other multifunctional RNA helicases, such as those involved in RNA processing and decay, are recruited to diverse substrates via dedicated cofactor proteins^74,75^, DHX15 and its associated G-patch proteins is the most extensive example of a single enzyme regulated by a network of different cofactors. The RNA/RNP remodeling force of a highly active, processive RNA helicase is required across gene expression processes, and the targeting of a single helicase possessing such properties to different processes via a network of related cofactors is not only an efficient mechanism via which to fulfil these individual functions but is also an elegant way in which different gene expression processes can be interconnected. Crosstalk between the different aspects of gene expression involving DHX15 and its G-patch protein cofactors, such as ribosome biogenesis, spliceosome disassembly/quality control and cap-proximal pre-mRNA methylation, is important for maintaining the correct balance between different RNPs involved in gene expression^40^. As the vast majority of DHX15–G-patch protein complexes function in the context of alternative splicing (**Fig. 6**; see for example^14,42,67,76,77^), cross-regulation across the G-patch protein network is a means to fine-tune the cellular transcriptome in different conditions. The expression levels of individual G-patch proteins vary considerably between cell types, and many G-patch proteins are differentially expressed across different tissues and in cancer^14,15,42,78–80,81^. Due to the mutually exclusive interactions of G-patch proteins with DHX15 and the ability of some G-patch proteins to outcompete each other for helicase association^82^, such alterations in the G-patch protein network will lead to re-distribution of the helicase between its different substrates.

Alongside their association with the multifunctional RNA helicase DHX15, an emerging characteristic of many G-patch proteins is their own multifunctionality, with several displaying both DHX15-associated and DHX15-independent functions. Our immunoprecipitation experiments demonstrate that only a small proportion of some G-patch proteins is associated with DHX15, while the vast majority remains unbound (**Fig. 1**). For example, while TFIP11 supports the function of DHX15 in spliceosome disassembly, a DHX15-independent function of the G-patch protein in facilitating association of snoRNPs with the U6 snRNA is described^53,83^. Likewise, GPATCH4 influences rRNA and snRNA modification via DHX15-associated and autonomous mechanisms^17^. Our findings highlight ZGPAT as another multifunctional G-patch protein; initially characterized as a DNA-binding transcription regulator^81^, ZGPAT was later identified a component of a stalled snRNP complex containing the U4 and U6 snRNAs (**Fig.5d, e**)^39^ and here, we reveal that this G-patch protein is required for efficient 2′-*O*-methylation of the U2 and U5 snRNAs (**Fig. 5i-k**). In contrast to other G-patch proteins that associate with introns containing sno/scaRNAs, ZGPAT interacts with excised scaRNAs (**Fig. 5b, f**). Although protein components of scaRNPs were not recovered with ZGPAT under the stringent conditions of the IP-MS experiments, the enrichment of ZGPAT in nuclear foci likely representing Cajal bodies (**Fig. 3**) strongly suggests association of this G-patch protein with scaRNPs. Depletion of ZGPAT alters 2′-*O*-methylation levels of specific nucleotides towards the 5′ end of the U2 snRNA and in the U5 snRNA. Dissection of the effects of snRNA modifications on pre-mRNA splicing has shown that the 2′-*O*-methylations within the first 20 nucleotides of U2 are dispensable for U2 snRNP assembly but are each individually required for correct spliceosomal E complex formation and efficient intron excision^84,85^. By contrast, the region of U5 containing the 2′-*O*-methylation impacted by depletion of ZGPAT (Um41) basepairs with the 5′ exon of the substrate pre-mRNA and is implicated in optimizing this interaction^86,87^. Our RIP-seq analyses indicate association of ZGPAT with only a minimal number of (pre-)mRNAs (**Fig. 5a**), suggesting that the transcriptome-wide alternative splicing changes observed upon depletion of ZGPAT (**Fig. 6**; **Supplementary Fig. S4**) likely arise due to perturbed snRNA 2′-*O*-methylation. The mechanistic basis of how ZGPAT influences efficiency of snRNA 2′-*O*-methylation remains unknown but the robust association of the G-patch protein with the scaRNAs guiding the affected methylations (**Fig. 5d**) suggests that it impacts either their assembly into functional methylation complexes or their association with snRNAs. Notably, ZGPAT also associates with other scaRNAs guiding snRNA modifications not observed to be affected by transient depletion of ZGPAT (**Fig. 5d, f**). It is possible, therefore, that the more widespread effects on snRNA modification may be observed upon more prolonged or complete depletion of ZGPAT. Similar to both GPATCH4 and TFIP11 that are required for efficient 2′-*O*-methylation^17,53^, ZGPAT likely influences snRNA modification independently of DHX15, as the helicase does not demonstrably associate with scaRNAs (**Fig. 5b, f**).

Most other surveyed G-patch protein RNA interactomes highlighted association with pre-mRNAs and pleiotropic effects of G-patch protein depletion were observed by our transcriptome-wide analyses (**Figs. 4b and 6**). A broad range of numbers of transcripts passed the high stringency thresholds in the RIP-seq experiments for each G-patch protein. It cannot be excluded that this arises due to differences in the modes of RNA association across the G-patch proteins, resulting in some interactomes being more of less comprehensively captured by the common approach applied. However, protein interactome experiments performed under similar conditions demonstrated robust association with spliceosomal complexes (**Fig. 4a**), suggesting that these are rather *bona fide* differences reflecting the functional relevance of individual G-patch proteins for diverse subsets of transcripts. Given the variable expression levels of the G-patch proteins, and the contribution of alternative splicing to dynamic gene expression reprograming, it is tempting to speculate that some G-patch proteins for which few RNA targets were recovered may become more functionally relevant in particular cellular conditions. For example, GPATCH3, which recovered minimal RNAs in RIP-seq experiments and whose depletion only nominally influences alternative splicing and gene expression, is implicated, with DHX15, in the innate immune response to viral infection^77,88^.

GPATCH8 and SUGP2 emerge, alongside DHX15, as G-patch proteins that each bind several hundreds of pre-mRNAs. While many RNA targets of these G-patch proteins are unique, they also associate with transcripts recovered with other G-patch proteins and thus their binding profiles are consistent with roles as global mRNA binding proteins. Despite the substantial overlap between transcripts bound by GPATCH8 and DHX15 (**Fig. 4c**; **Supplementary Fig. S2b**) and the impact of depletion of the helicase or G-patch protein on splicing of these pre-mRNAs, the directionality of AS changes is generally different between DHX15 and GPATCH8. This is notably different to other G-patch proteins, for example, SUGP1, that not only associates with a subset of transcripts shared with DHX15, but also similarly influences splicing and expression of these transcripts. Consistent with these data, SUGP1 was recently characterized as supporting splicing fidelity control by DHX15 and functionally opposed to GPATCH8^14,67^. These data further support the concept of DHX15-associated and DHX15-independent functions of G-patch proteins as well as the notion that interplays between different members of the G-patch protein family can fine-tune gene expression.

Despite the recent insights into the role of SUGP1 in splicing regulation^14,67,89^, the related SUGP2 has remained poorly characterized functionally. Both proteins contain tandem SURP domains together with a C-terminal G-patch, but SUGP1 also includes a short U2AF ligand motif (ULM) that contributes to interaction with DHX15^67^. As this feature is lacking from SUGP2, this may, at least in part, rationalize the weak recovery of DHX15 with SUGP2 (**Fig. 1c**) and the functional divergence of these related proteins. While SUGP1– DHX15 is implicated in repressing neo-junction splicing and binds to introns near branch points^14^, our data show a broader distribution of intron binding for SUGP2 (**Fig. 7**) that is reminiscent of other deep-intron binding RBPs. In this context, SUGP2 was described, among 28 RBPs, as LINE element-binding^90,91^. Most LINEs within the genome are degenerate and thus not active retrotransposable elements^92^, but nevertheless harbor many potential splice sites and play roles in tissue-specific exon splicing^91^. Other deep intron-binding proteins, such as MATR3, PTBP1 and hnRNPM were shown to repress splice and poly(A) sites. The reduced splicing efficiency of introns bound by SUGP2, together with increased splicing and expression upon SUGP2 depletion (**Fig. 6g, Fig. 7a-d**) supports the hypothesis that intron-binding by SUGP2 masks spliceosome recognition features to suppress splicing. Interestingly, MATR3, PTBP1 and hnRNPM as well as various other hnRNPs were co-precipitated with SUGP2 (**Fig. 4a**), suggesting that the G-patch proteins functions synergistically with other splicing repressors. As pre-mRNA splicing is initiated co-transcriptionally^93,94^, a role of SUGP2 in splicing suppression necessitates its association with nascent RNAs. The enrichment of SUGP2 in chromatin as well as the finding that it binds pre-mRNAs and lncRNAs encoded in chromosomal proximity support that SUGP2 does indeed bind transcripts co-transcriptionally (**Fig. 7g-i**). Cryptic splice sites and pseudo exons are prevalent features of introns that are often implicated in genetic diseases^95,96^. Hence, the identification of SUGP2 as a splicing repressor highlights an important mechanism of gene expression regulation by this G-patch protein.

Considered collectively, the family of G-patch proteins comes to the fore as a diverse group of proteins that function both as RNA helicase cofactors and independently to influence many different aspects of gene expression. These systematic analyses of the human G-patch protein family reveal fundamental principles of G-patch protein-mediated helicase regulation and common targets of G-patch proteins and their cognate RNA helicases as well as unique features and individual functions of G-patch proteins. Focused analyses of specific G-patch proteins highlight distinct modes of pre-mRNA splicing regulation, including indirect regulation via modulation of snRNA modifications, helicase-associated alternative splicing modulation and splicing suppression by deep intron binding. However, extensive crosstalk has also become evident at the levels of expression regulation and functional activity, both between G-patch proteins and their respective RNA helicase partners as well as across the G-patch protein network.

## MATERIALS AND METHODS

### Molecular cloning

Standard molecular biology methods^97^ were used to generate the plasmids listed in **Supplementary Table S1**. The coding sequences (CDS) of the human G-patch proteins (AGGF1 - NM_018046, CHERP - NM_006387, CMTR1 - NM_015050, GPANK1 - NM_001199237, GPATCH1 - NM_018025, GPATCH2 - NM_018040, GATCH3 - NM_022078, GPATCH8 - NM_001002909, GPATCH11 - NM_174931.4, GPKOW - NM_015698, PINX1 - NM_017884, RBM5 - NM_005778, RBM6 - NM_005777, RBM10 - NM_005676, RBM17 - NM_001145547, SUGP1 - NM_172231, SUGP2 - NM_001017392, TFIP11 - NM_001008697, ZGPAT - NM_001195653 as well as the RNA helicase DHX35 (NM_021931) were cloned into pcDNA5-based plasmids for the inducible expression of C-terminally His_6_-2xFLAG-tagged or N-terminally 2xFLAG-His_6_-tagged (FLAG-tagged) proteins in human HEK293 Flp-In T-REx cells (Thermo Fisher Scientific). Site-directed mutagenesis was performed to induce the following amino acid exchanges in the expressed proteins: GPATCH1 glycine 156 to glutamate (G156E) and DHX35 leucine 460 to glutamate (L460E).

For recombinant expression of proteins/peptides in *Escherichia coli* (*E. coli*), the sequences encoding the G-patch motifs of the human G-patch proteins, as defined by Uniprot, and the CDS of GPKOW were cloned into pQE80-derived vectors for the expression of N-terminally ZZ and C-terminally His_7_-proteins (**Supplementary Table S1**). The CDSs of DHX16 (NM_003587) and DHX35 were cloned into pQE80-derived vectors for the expression of N-terminally maltose binding protein (MBP) and C-terminally His_10_-tagged proteins. A plasmid for the expression of DHX15 with equivalent tags was previously described^16^.

### Human cell culture and RNAi

HEK293 Flp-In T-REx cells were maintained in DMEM containing 10% fetal bovine serum (FBS) and 1% penicillin-streptomycin under standard conditions (37°C and 5% CO_2_). Cell lines used in this study were routinely checked for mycoplasma contamination with the *Mycoplasmacheck* service (Eurofins Genomics). For generation of stably transfected cell lines, HEK293 Flp-In T-REx cells were co-transfected with the pOG44 plasmid encoding the Flp recombinase and the appropriate pcDNA5/FRT/TO-derived plasmid (**Supplementary Table S1**) using the X-tremeGENE 9 DNA transfection reagent (Roche) according to the manufacturer’s instructions. Cells with correct *trans*gene insertion were selected using 82.4 µg/ml Hygromycin B (Applichem) and 10 µg/ml Blasticidin S Hydrochloride (Applichem). To induce expression of the encoded proteins, cells were treated for 24 h with 1 µg/ml tetracycline.

RNAi-mediated depletion of proteins from HEK293 Flp-In T-REx cells was performed by transfecting 20-50 nM siRNAs (**Supplementary Table S2**) with Lipofectamine RNAiMAX (Thermo Fisher Scientific) following manufacturer’s protocol for reverse transfections. Cells were harvested 72-96 h after transfection.

### RNA extraction, cDNA synthesis, qPCR and RT-PCR

Total RNA was extracted from human cells using TRI Reagent (Sigma Aldrich) according to the manufacturer’s instructions. First-strand cDNA synthesis was performed using the SuperScript III Reverse Transcriptase kit (Thermo Fisher) according to the manufacturer’s instructions using 2 µg total RNA and random hexamer primers (75 pmol final concentration; Sigma Aldrich). Quantitative RT-PCR was performed in a LightCycler 480 (Roche) using the LightCycler 480 SYBR Green I Master kit (Roche). Each reaction contained 0.65x SYBR Green mix, 0.33 pmol each forward and reverse primer (**Supplementary Table S3**) and 3 µl cDNA (diluted 1:10). Amplification was performed using the following program: 5 min at 95 °C and 50 cycles of 10 sec denaturation at 95 °C, 20 sec annealing at 58 °C and 15 sec elongation at 72 °C. Melting curve analysis was subsequently performed by incubation for 10 sec at 95 °C and 1 min at 55 °C followed by continuous acquisition of fluorescence until 97 °C. All primers were verified for efficient amplification prior to use and generated a single product as determined by melting curve analysis. Technical triplicates were pipetted for each biological replicate. Data analysis steps were performed using the LightCycler 480 software and results were normalized to the expression level of the *EMC7* mRNA, which is a verified housekeeping gene recommended for qPCR relative quantification^98^. For RT-PCRs monitoring alternative splicing, PCR reactions (26 cycles) were carried out using Phusion polymerase and primers listed in **Supplementary Table S3**. Reaction products were separated on 4% polyacrylamide gels in 1x TBE and detected via SYBRGold staining using a Typhoon FLA 9500 scanner (Cytiva).

### Immunofluorescence

Stably transfected HEK293 Flp-In T-REx-derived cell lines were grown on sterile glass coverslips coated with 0.01% poly-L-lysine (Sigma-Aldrich) and expression of *trans*genes was induced for 24 h with 1 µg/ml tetracycline. Cells were fixed with 2.5% paraformaldehyde in PBS for 15 min and then washed two times with phosphate-buffered saline (PBS) for 5 min. Permeabilization was done for 20 min with 0.1% Triton X-100 in PBS and then, cells were blocked for 1.5 h with PBS containing 1% FBS and 0.1% Triton X-100. Cells were incubated with the primary antibodies (**Supplementary Table S4**) for 2 h, washed three times for 10 min each with PBS and stained for 1.5 h with Alexa Fluor 488 and 594-conjugated secondary antibodies (**Supplementary Table S4**). The cells were then washed with PBS and mounted on microscope slides using VECTASHIELD Antifade Mounting Medium with DAPI (Vector Labs). Cells were imaged using a ZEISS LSM 510 META laser scanning microscope with a 100x objective and images were processed using Fiji.

### Immunoprecipitation of complexes containing FLAG-tagged proteins

HEK293 Flp-In T-REx cell lines expressing FLAG-tagged proteins resuspended in IP buffer (20 mM HEPES-NaOH pH 7.5, 250 mM NaCl, 0.5% Triton X-100, 10% glycerol, 0.5 mM EDTA) supplemented with cOmplete Mini Protease Inhibitor Cocktail (Roche) were lysed by sonication (20% amplitude, 3 cycles of 15 sec with 0.3 sec on/0.7 off) using a Branson Digital Sonifier. Cell debris were pelleted by centrifugation for 10 min at 20,000 g and 4 °C and the soluble lysate was added to 30 µl of anti-FLAG M2 Magnetic Beads (Sigma-Aldrich) that had been pre-equilibrated in IP buffer. Binding was carried out for 2 h at 4 °C in the presence of 50 µg/ml RNase A (Applichem). The beads were then washed five times with the IP buffer and bound complexes were eluted for 30 min at 4 °C using 250 µg/ml FLAG Peptide (Sigma-Aldrich) diluted in IP buffer. Proteins in the eluates were precipitated with 20% trichloroacetic acid for 20 min on ice and pelleted by centrifugation for 20 min at 20,000 g and 4 °C. Protein pellets were washed with ice-cold acetone and air-dried briefly. For mass spectrometry analysis, samples were resuspended in 1x NuPAGE LDS Sample Buffer (Thermo Fisher Scientific) supplemented with 50 mM DTT and denatured at 70 °C for 10 min. For western blotting analysis, the pellets were resuspended first in 100 mM Tris-HCl pH 8.4, mixed with 4x SDS loading dye (200 mM Tris-HCl pH 6.8, 8% SDS, 50 mM EDTA, 4% 2-mercaptoethanol, 0.08% bromophenol blue, 40% glycerol) and incubated at 95 °C for 10 min.

### Liquid chromatography tandem mass spectrometry (LC-MS/MS) of IP eluates

Samples were reconstituted in 1× NuPAGE LDS Sample Buffer (Invitrogen) and separated on 4-12 % NuPAGE Novex Bis-Tris Minigels (Invitrogen). Gels were stained with Coomassie Blue, and each lane sliced into 12 equidistant gel slices regardless of staining. After washing, slices were reduced with dithiothreitol (DTT), alkylated with 2-iodoacetamide and digested with trypsin overnight. The resulting peptide mixtures were then extracted, dried in a SpeedVac, reconstituted in 2% acetonitrile/0.1% formic acid/ (v:v) and prepared for nanoLC-MS/MS as described previously^99^.

For mass spectrometric analysis samples were enriched on a self-packed reversed phase-C18 precolumn (0.15 mm ID x 20 mm, Reprosil-Pur120 C18-AQ 5 µm, Dr. Maisch, Ammerbuch-Entringen, Germany) and separated on an analytical reversed phase-C18 column (0.075 mm ID x 200 mm, Reprosil-Pur 120 C18-AQ, 3 µm, Dr. Maisch) using a 30 min linear gradient of 5-35 % acetonitrile/0.1% formic acid (v:v) at 300 nl min-1). The eluent was analyzed on a Q Exactive hybrid quadrupole/orbitrap mass spectrometer (ThermoFisher Scientific, Dreieich, Germany) equipped with a FlexIon nanoSpray source and operated under Excalibur 2.4 software using a data-dependent acquisition method. Each experimental cycle was of the following form: one full MS scan across the 350-1600 *m/z* range was acquired at a resolution setting of 70,000 FWHM, and AGC target of 1*10e6 and a maximum fill time of 60 ms. Up to the 12 most abundant peptide precursors of charge states 2 to 5 above a 2*10e4 intensity threshold were then sequentially isolated at 2.0 FWHM isolation width, fragmented with nitrogen at a normalized collision energy setting of 25%, and the resulting product ion spectra recorded at a resolution setting of 17,500 FWHM, and AGC target of 2*10e5 and a maximum fill time of 60 ms. Selected precursor *m/z* values were then excluded for the following 15 s. Two technical replicates per sample were acquired.

### LC-MS/MS data analysis

Peaklists were extracted from the raw data using Raw2MSMS software v1.17 (Max Planck Institute for Biochemistry, Martinsried, Germany), and analyzed using Mascot (Matrix Science, London, UK; version 2.5.1). Mascot was set up to search the human SwissProt_2017_04 reference proteome (20199 entries) assuming the digestion enzyme trypsin. Mascot was searched with a fragment ion mass tolerance of 0.050 Da and a parent ion tolerance of 10.0 ppm. Carbamidomethyl of cysteine was specified in Mascot as a fixed modification. Oxidation of methionine was specified in Mascot as a variable modification.

Scaffold (version Scaffold_5.3.0, Proteome Software Inc., Portland, OR) was used to validate MS/MS based peptide and protein identifications. Peptide identifications were accepted if they could be established at greater than 91.0% probability to achieve an FDR less than 1.0% by the Scaffold Local FDR algorithm. Protein identifications were accepted if they could be established at greater than 6.0% probability to achieve an FDR less than 1.0% and contained at least 2 identified peptides. Protein probabilities were assigned by the Protein Prophet algorithm^100^. Proteins that contained similar peptides and could not be differentiated based on MS/MS analysis alone were grouped to satisfy the principles of parsimony. Proteins sharing significant peptide evidence were grouped into clusters. Only proteins with total spectral counts ≥10 and an enrichment of at least 1.5-fold with the FLAG-tagged G-patch protein compared to the control (FLAG tag only) were considered as enriched for further analysis.

### Western blotting

Following separation by SDS-polyacrylamide gel electrophoresis (SDS-PAGE), proteins were transferred onto Hybond P 0.45 PVDF or nitrocellulose blotting membranes (Cytiva) using a wet-transfer system. Membranes were blocked for 1 h at room temperature using 5% milk in TBS with 0.1% Tween-20 (TBS-T), followed by incubation with primary antibodies (**Supplementary Table S4**) overnight at 4°C. Membranes were washed three times for 10 min in TBS-T and incubated with HRP-coupled/IRDye-conjugated secondary antibodies (**Supplementary Table S4**) for 1 h at room temperature. After washing with TBS-T, signals were detected using the Immobilon Western Chemiluminescent HRP Substrate (Merck) or scanned using an Odyssey CLx scanner (LI-COR Biosciences).

### Recombinant protein expression in *E. coli* and protein purification

Plasmids for the recombinant expression of proteins in *E. coli* (**Supplementary Table S1**) were used to transform BL21 (DE3) pLysS (MBP-DHX15-His, MBP-DHX16-His, MBP-DHX35-His) or BL21 (DE3) CodonPlus-RIL cells (ZZ-G-patch-His, ZZ-GPKOW_FL_-His)). Induction of protein expression in liquid cultures was done either with 500 mM isopropyl-β-D-thiogalactopyranoside (IPTG) for 3 h at 37 °C for the G-patch motifs or with 250 mM IPTG at 18 °C overnight in the case of DHX15, DHX16, DHX35 and GPKOW_FL_. Cells were harvested by centrifugation at 5000 g for 10 min and all subsequent protein purification steps were done at 4 °C.

For purification of ZZ-His-tagged G-patch motifs, cell pellets were resuspended in lysis buffer (50 mM Tris-HCl pH 7.4, 150 mM NaCl, 1 mM MgCl_2_, 10% glycerol, 5 mM imidazole) and sonicated on ice using a Branson Digital Sonifier at 45% amplitude for 4 cycles with 0.5 sec on/0.5 sec off pulses and with 30 sec pause between the cycles. The lysate was cleared by centrifugation at 20,000 g for 20 min and His-tagged proteins were retrieved by 1 h incubation with 1 ml Ni-NTA resin (Roche) that had been pre-equilibrated in the lysis buffer. The beads were washed thoroughly with wash buffer 1 (50 mM Tris-HCl pH 7.4, 150 mM NaCl, 1 mM MgCl_2_, 10 mM imidazole), then wash buffer 2 (50 mM Tris-HCl pH 7.4, 1 M NaCl, 1 mM MgCl_2_, 10 mM imidazole) and again with wash buffer 1. Elution was carried out using 50 mM Tris-HCl pH 7.4, 150 mM NaCl, 1 mM MgCl_2_, 250 mM imidazole. Fractions containing the highest amounts of protein were pooled and dialyzed against 50 mM Tris-HCl pH 7.4, 150 mM NaCl, 1 mM MgCl_2_ and 10% glycerol.

For purification of MBP-DHX15-His, MBP-DHX16-His, MBP-DHX35-His and ZZ-GPKOW_FL_-His, cells were resuspended in 50 mM Tris-HCl pH 7.4, 600 mM NaCl, 1 mM MgCl_2_, 0.5% Triton X-100, 10% glycerol, 25 mM imidazole, 2 mM 2-mercaptoethanol and lysed with EmulsiFlex-C3 (Avestin) by three passes at 10,000 psi. The cleared lysate was incubated with pre-equilibrated Ni-NTA beads (Qiagen) for 1.5 h. The beads were then washed with 50 mM Tris-HCl pH 7.4, 150 mM NaCl, 1 mM MgCl_2_, 10% glycerol, 40 mM imidazole followed by a wash step with the same buffer containing 1 M NaCl before a final washing step with the initial wash buffer. Bound proteins were eluted using 50 mM Tris-HCl pH 7.4, 150 mM NaCl, 1 mM MgCl_2_, 10% glycerol, 250 mM imidazole and the fractions containing the highest amount of protein were pooled. Buffer exchange to 50 mM Tris-HCl pH 7.4, 150 mM NaCl, 1.5 mM MgCl_2_ and 10% glycerol was done on PD-10 columns (Cytiva). Protein concentrations were determined using the Coomassie Plus (Bradford) assay (Thermo Fisher) and proteins were visualized by SDS-PAGE followed by Comassie staining (0.1% Coomassie R-250, 10% acetic acid, 40% methanol) followed by destaining with a solution of 10% acetic acid and 20% methanol.

### Structure modeling

The AlphaFold server (https://alphafoldserver.com/) was used to generate a model of GPATCH1 (Q9BRR8), DHX35 (Q9H5Z1) and WDR83 (Q9BRX9) as well as DHX15 (Q8IX01) with GPATCH8 (Q9UKJ3) or SUGP2 (Q8IX01). Predicted aligned error (PAE) scores were plotted using PAE Viewer (https://pae-viewer.uni-goettingen.de/).

### NADH-coupled ATPase assays

Hydrolysis of ATPase by RNA helicases was monitored using an NADH-coupled enzymatic assay^101^. Reactions containing 45 mM Tris-HCl pH 7.4, 25 mM NaCl, 2.5 mM MgCl_2_, 1 mM phosphoenolpyruvate, 300 µM NADH, 20 U/ml pyruvate kinase/lactic dehydrogenase, 4 mM ATP and 2 µM U_32_ RNA oligonucleotide (IDT) were set up and MBP-DHX15-His, MBP-DHX16-His or MBP-DHX35-His were added to final concentrations of 250 nM. Reactions were further supplemented with 1.5 µM ZZ- and His-tagged G-patch domains as required. Experiments were carried out at 30°C and the absorbance at 340 nm was measured every min for 50 min using a BioTEK Synergy plate reader. The amount of ATP hydrolyzed, which is equimolar to the amount of NADH oxidized, was determined from the slope of the linear absorbance decrease.

### Fluorescence anisotropy

Recombinant proteins were re-dialyzed for 16 h against a buffer containing 50 mM Tris-HCl pH 7.4, 90 mM NaCl, 1 mM MgCl_2_, 4% glycerol. Anisotropy of 50 nM fluorescently-labeled RNA (5′-GUAAUGAAAGU-ATTO647-3′) was measured in the presence of 0-600 nM MBP-DHX15-His and the reactions were supplemented with the ZZ-/His-tagged G-patch motifs or the ZZ-His tags only at a concentration of 1.2 µM. Alternatively, to determine the RNA affinities of the G-patch motifs of GPATCH1, GPATCH2, GPATCH8 and GPKOW, protein concentrations of 0-10 µM were analyzed. Anisotropy measurements were performed at room temperature with a FluoroMax-4 spectrofluorometer (Horiba Scientific) using the following settings: excitation wavelength - 644 nm, emission wavelength - 661 nm, excitation and emission slits - 8, integration time - 0.5 sec, temperature - 25°C. Data were analyzed using the following equation:

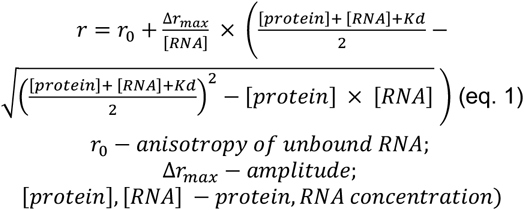

For reactions containing both DHX15 and G-patch cofactors, only the concentration of the helicase was considered for calculations due to the insignificant level of RNA binding displayed by the G-patch motifs themselves.

### Crosslinked RNA immunoprecipitation (RIP)-seq

HEK293 Flp-In T-REx cell lines expressing FLAG-tagged proteins were crosslinked using UV light at 254 nm with 400 mJ/cm^2^ in a Stratalinker UV Crosslinker (Stratagene). The harvested cells were resuspended in 1 ml RIP buffer (20 mM HEPES-NaOH pH 7.5, 250 mM NaCl, 1 mM MgCl_2_, 0.5% Triton X-100, 10% glycerol) containing cOmplete Mini Protease Inhibitor Cocktail (Roche) and 80 U RNasin Ribonuclease Inhibitor (Promega) then lysed by sonication with a Branson Digital Sonifier at 25% amplitude in 3 cycles of 15 sec (0.3 sec on/0.7 sec off) with 30 sec pause in between. After centrifugation for 15 min at 20,000 g and 4 °C, the cleared cell lysate was incubated for 2 h with 50 µl of anti-FLAG M2 Magnetic Beads (Sigma-Aldrich) that had been pre-equilibrated in RIP buffer. The beads were washed five times with RIP buffer and bound complexes were eluted by incubation with 250 µg/ml FLAG Peptide (Sigma-Aldrich) diluted in RIP buffer for 2 h at 4 °C. The eluates were treated with 2 U TURBO DNase (Thermo Fisher) for 20 min at 37°C in the presence of 40 U RNasin Ribonuclease Inhibitor (Promega) before treatment with 275 µg/ml Proteinase K (Roche) for 2 h at 42 °C in a reaction containing 1% SDS and 0.5 mM EDTA. RNA was extracted with an equal volume of phenol-chloroform-isoamyl alcohol (25:24:1) in the presence of 0.3 M sodium acetate pH 5.2. before ethanol precipitation. RNA pellets were washed twice with 75% ethanol and resuspended in nuclease-free water (Qiagen). RNA samples were subjected to ribosomal RNA (rRNA) depletion and then converted into cDNA libraries using the Illumina TruSeq Stranded Total RNA Kit. Single-end sequencing of the libraries was done with a read length of 50 bp on an Illumina HiSeq 4000 sequencer and generated approximately 20-50 million reads per sample. Library preparation and sequencing were performed by the NGS Integrative Genomics Core Unit (University Medical Center Göttingen), followed by bioinformatic analysis described below. Alternatively, RNAs in inputs and eluates were analyzed by northern blotting.

### Northern blotting

Total RNA samples or RNAs extracted from input and eluate samples of RNA-IP experiments were separated on denaturing (7 M urea) 8-12% polyacrylamide gels in TBE (90 mM Tris-HCl, 90 mM boric acid, 2.55 mM EDTA) and transferred to Hybond-N membrane (Cytiva) in 0.5x TBE. RNAs were crosslinked to membranes using 240 mJ/cm^2^ light at 254 nm in a Stratalinker UV crosslinker (Stratagene). Membranes were pre-hybridized in SES1 buffer (250 mM sodium phosphate pH 7.0, 7% SDS, 1 mM EDTA) for 40 min at 37 °C and 5′ [^32^P]-labelled antisense DNA oligonucleotide probes (**Supplementary Table S5**), diluted in SES1 buffer were added overnight at 37 °C to detect the RNA species of interest. Membranes were washed sequentially with 6x SSC (150 mM NaCl, 15 mM sodium citrate) and 2x SSC supplemented with 0.1% SDS for 30 min each at 37 °C. After drying, membranes were exposed to a phosphorimager screen and signals were detected with a Typhoon FLA9500 scanner (Cytiva). Images were processed using ImageQuant software (Cytiva).

### RiboMeth-seq

For RiboMeth-seq^59,102^, 5-10 μg nuclear RNA from HeLa cells treated with non-target siRNAs or those targeting ZGPAT, DHX15, or both, was partially degraded by alkaline at denaturing temperature. The RNA fragments were separated on a denaturing (8 M urea) polyacrylamide gel, and fragments in the size range of 20–40 nucleotides were excised and ligated to adapters using a modified Arabidopsis thaliana tRNA ligase to join 2′-3′ cyclic phosphates and 5′ phosphorylated ends. cDNA was generated using Superscript III reverse transcriptase and libraries were sequenced on the Ion Proton platform. Reads were mapped to the human snRNA sequences (genome build hg38) using Bowtie2 (v. 2.3.4.1), and the RMS score (fraction methylated) was calculated as ‘score C’ in^103^, expect for cap-proximal methylations in snRNAs that was calculated according to^104^. In a few cases, a barcode correction was applied when calculating the RMS score as described previously.

### RNA-seq

HEK293 Flp-In T-REx cells were treated with the indicated siRNAs and total RNA was extracted using TRI Reagent (Sigma-Aldrich) according to the manufacturer’s instructions. RNAs were incubated with 2 U TURBO DNase (Thermo Fisher) for 15 min at 37°C in the presence of 40 U RNasin Ribonuclease Inhibitor (Promega) before re-purification using the RNeasy Mini Kit (Qiagen) according to manufacturer’s instructions. Total RNA samples were depleted of rRNAs and converted into cDNA libraries using the Illumina TruSeq Stranded Total RNA Kit (NGS Integrative Genomics Core Unit (University Medical Center Göttingen)). Single-end sequencing of the libraries was carried out on an Illumina HiSeq 4000 sequencer, and bioinformatic analyses were performed as described below.

### RIP-seq and RNA-seq mapping

FASTQC 0.11.9 from the FASTX toolkit was run to ensure high quality of the data before mapping (http://hannonlab.cshl.edu/fastx_toolkit/index.html). For mapping, the aligner STAR 2.7.11b with the following settings was used: --outFilterMultimapNmax 10 -- outSAMattributes All --outSAMtype BAM SortedByCoordinate --outReadsUnmapped Fastx -- chimSegmentMin 20 --chimOutType WithinBAM Junctions --quantMode TranscriptomeSAM GeneCounts --outWigType bedGraph --outWigNorm RPM --outWigStrand Unstranded -- outFilterMismatchNmax 2. Bedgraph files were generated using BEDTools (v2.27.1). UCSC Genome Browser tools bedSort (v469) and bedGraphToBigWig (v469) were used to convert from bedgraph format to bigwig format for visualization of the data in the IGV genome browser. To index and view bam files samtools 1.21 was used. Genome assembly version GRCh38.p14 and the GENCODE gene set from release 47 were used and gene, exon, intron and transcript annotations from ensembl, entrez and hgnc were obtained with the Bioconductor package biomaRt.

### Sashimi plots

Individual RIP- or RNA-seq replicates were pooled using samtools 1.21. Visualization was done using ggsashimi v1.1.5^105^ with a minimum junction coverage of 2 for RIP samples and 4 for RNA-seq samples.

### RNA abundance quantification in RIP-seq

RNA abundance was quantified using the DESeqDataSetFromHTSeqCount, DESeq and results functions from the DESeq2 package in Bioconductor^106^ for RIP experiments with multiple replicates (DHX15, DHX35, GPATCH1, RBM17). Read counts per gene were obtained using bedtools intersect from the BEDTools (v2.27.1) suite and only reads spanning less distance than the gene length were counted. Genes with 10 or more reads in at least one replicate were kept for initial abundance quantification. For RIP-seq samples with only one replicate at least 10 reads in the IP or background control sample were required for further analysis. Read counts across all genes were median normalized for the IP and background control sample, respectively. To estimate enrichment of RNAs upon immunoprecipitation, log2-fold changes between IP and background control were calculated and a pseudo-count of 0.001 was introduced to include genes that had no detectable signal in the background, but in the IP.

### Defining RIP-seq targets

#### Lower and upper bound as cutoff

For defining RIP hits, namely RNAs that are bound/crosslinked to the bait protein and thus detected in RIP-seq, a cutoff needed to be defined above which genes could be considered enriched in the RIP sample. Based on random noise in the data, log2-fold changes varied by several fold. We expected a roughly symmetrical distribution if there would be no RIP hits, whereas a bait with many RNA binding targets would result in a highly skewed, i.e. right-tailed, distribution (**Supplemental Fig. S2a**). We assumed that ‘derichment’, i.e. negative log2-fold changes relative to the distribution median, originated largely from random variation and noise in the experiment and could be used to identify ‘true’ RIP hits in the right tail of the distribution. To this end, we defined a lower boundary (and corresponding symmetrical upper boundary) at which at most 5% of the gene counts with log2-fold changes greater than the upper bound can be explained by random variation.

To find the optimal lower bound, we probed lower quantile ranges from 0 to 5% in 0.2% intervals. We determined the log2-fold change at the lower quantile ranges (potential lower bound) and determined the corresponding upper bound, namely the log2-fold change at which the corresponding quantile boundary would be in a symmetrical distribution. This was determined as two times the distribution median minus the lower bound. Next, we assessed how many genes fall below or above the lower and upper bound, respectively. The final lower bound for individual baits was selected as the bound where the number of genes below the lower bound makes up at most 5% of the genes above the upper bound (approximately an FDR < 0.05). In cases where multiple lower quantile ranges fulfilled this criterion, the greatest lower quantile range was used. In cases where no lower quantile range could be identified with an FDR < 0.05, the lower quantile range with the next lowest FDR was used for bound RNA classification.

Additionally, genes with log2-fold changes greater than the upper quantile boundary were also required to have 50 reads or more in the RIP-seq sample and rpkms > 0 in a matching total RNA-seq control sample (described below in the siRNA-seq section) to be considered as RIP hits. Furthermore, for RIP samples analyzed with DEseq2 additional statistical confidence was gained by requiring an FDR < 0.05.

#### Comparison to background control replicates

The filtering criteria described above provided a reasonable selection of potential RIP hits, however some samples showed spurious hits like rRNA and mitochondrial rRNA, e.g. in the TFIP11 RIP-seq, that could be easily explained by less rRNA and mitochondrial rRNA signal in the matched background control than in typical other controls. It is reasonable to expect that RIP hits might be substantially biased by the inclusion of only one background control sample. Hence, multiple background control samples were prepared and could be used to assess variation in enrichment analysis based on variation in background RNA binding. To this end, log2-fold changes in read counts per gene were calculated for all combinations of RIP samples to the 7 background controls. Genes that were considered possible RIP hits from the analysis above between experimentally matched RIP sample and background control, were only considered further, if it was enriched relative to at least 50% of the other background samples. This refinement eliminated the spurious hits like rRNA and mitochondrial rRNA and thus resulted in a refined list of possible target RNAs of RIP baits. Median log2-fold changes of RIP / background controls are used for visualization and correlation between conditions.

### RNA-seq expression analysis

As for RNA abundance quantification in RIP-seq, read counts per gene were obtained using bedtools intersect from the BEDTools (v2.27.1) suite and only reads spanning less distance than the gene length were counted. Gene expression was quantified using the DESeqDataSetFromMatrix, DESeq and results functions from the DESeq2 package in Bioconductor^106^. Three biological replicates of any G-patch protein or RNA helicase knock-down were used for analysis with respectively matched non-target siRNA controls. Individual replicates were prepared in different batches; hence batch and condition were specified in the design flag of the DESeqDataSetFromMatrix function. The DeSeq function was applied for genes that had 50 or more reads in at least one sample. Differential expression was quantified for every knock-down relative to control separately. Hence, the ‘baseMean’ expression value captures the average expression for that gene in the respective knock-down and control. DESeq2 output that was used for further analysis in **Fig. 6, 7, Supplementary Figs. S4 and S5** can be found in **Supplementary Data 7**.

### RNA-seq alternative splicing analysis

rMATS^107^ was used for the analysis of alternative splicing (AS) patterns of paired replicates. To this end rMATS version 4.1.2 was used in combination with R version 3.6.3: rmats.py --b1 <path to knock-down bam files> --b2 <path to siNT bam files> --gtf annotations/hg38/gencode.v47.chr_patch_hapl_ scaff.annotation.gtf --od out --tmp out_prep -t single -- readLength 50 --variable-read-length --libType fr-firststrand --task both --paired-stats --mel 500 --allow-clipping.

Only AS events with a median of 5 junction reads or more in the 3 replicates of either skipped or included junctions per condition (SJC1, SJC2, IJC1, IJC2) were included for further analysis and the median inclusion level difference computed in addition to the rMATS output of the mean inclusion level difference. The median inclusion level difference distribution was used for the analysis presented in **Fig. 6b, d, f, Supplementary Fig. S4c, d** with a significance cutoff of p-Value < 0.05. rMATS output can be found in **Supplementary Data 7**.

With the default setting of mel 500 only short & annotated as retained introns are included. To include more introns, a previously used strategy^62^ of counting spliced and unspliced junction reads at annotated introns and defining IR as 1 - splicing efficiency, similar to FRASER^60^ or SPI^61^ was used. A minimal count of 20 junction reads was required for IR calculation. IR output can be found in **Supplementary Data 7**.

For both rMATS output and IR calculation, only events that passed all applied cutoffs in 80% of the datasets were considered for further analysis.

### SUGP2-related analysis

BigWig files for the two replicates of SUGP2 eCLIP in the HepG2 cell line were retrieved from the ENCODE platform^64^. Repeat annotations were retrieved from the UCSC tables browser (RepeatMasker Viz. track for hg38 genome annotation). Annotated regions of LINE1 (Long interspersed nuclear element-1) elements are displayed as collapsed tracks in **Fig. 7a, h**.

To quantify the fraction of intronic and exonic signal for different gene groups bedgraph coverage was intersected with intron and exon annotation obtained from biomart in Bioconductor / R using ENSEMBL_MART_ENSEMBL and hsapiens_gene_ensembl. The signal over all unique exons / introns of a given gene was summed and displayed as stacked barplot in **Fig. 7b**. The error bars reflect the standard deviation among different RIP-seq FLAG controls (n = 6) and the standard deviation between different siNT RNA-seq datasets (n = 8). The SUGP2 bound gene group refers to genes that were identified as enriched by SUGP2 RIP-seq (see above, **Fig. 4b**), random sampling of the same number of genes as in the SUGP2 bound group from annotated genes defines the random gene group.

For intron feature analysis intron length, nucleotide composition and splice site strength and PY-tract strength were considered (**Fig. 7, Supplementary Fig. S5**). The analysis was done as described previously^62,108^.

To probe for the genomic distance of genes encoding SUGP2 bound RNAs we assigned a rank / index to each gene by chromosome based on the mid-position of the gene. An index starting from 1 reflects the gene with the smallest mid-gene position on the respective chromosome. The difference between indices to the next neighbor for SUGP2 bound RNAs ordered by chromosome and mid-gene position was calculated and compared to 50 times randomly sampled gene groups of the same size as the SUGP2 bound RNA counts. Only non-overlapping genes were considered for this analysis. The Kolmogorov-Smirnov test with Benjamini & Hochberg correction was performed on every combination of SUGP2 bound RNAs to the random gene groups. The mean adjusted p-Value is given in **Fig. 7i**.

### Sub-cellular fractionation and analysis of chromatin-associated proteins

To prepare a nucleoplasmic extract and enrich chromatin-associated proteins^109^, 10 million cells were washed twice with ice-cold PBS and pelleted. 400 μL of Cell Lysis Buffer (10 mM Tris-HCl pH 7.4, 150 mM NaCl, 0.15% (v/v) NP-40, 1x protease inhibitors) was added to the pellet and cells were lysed by gentle pipetting followed by 5 min incubation on ice. The cell lysate was carefully overlaid onto 1 mL cold Sucrose Buffer (10 mM Tris-HCl pH 7.4, 150 mM NaCl, 24% (w/v) sucrose, 1x protease inhibitors) and centrifuged at 2,000 g, 4°C for 10 min to separate nuclei from the cytoplasmic fraction. The cytoplasmic supernatant was discarded and the nuclei were ruptured by resuspension in 250 μL of Glycerol Buffer (20 mM Tris-HCl pH 7.4, 75 mM NaCl, 0.5 mM EDTA, 50 % (v/v) glycerol, 1x protease/phosphatase inhibitors), followed by the immediate addition of 250 μL of Nuclear Lysis Buffer (10 mM Tris-HCl pH 7.4, 1 M urea, 300 mM NaCl, 7.5 mM MgCl_2_, 0.2 mM EDTA, 1% (v/v) NP-40, 1x protease inhibitors) and incubation on ice for 2 min. Chromatin was separated from the nucleoplasm by centrifugation at 13,000 g, 4°C for 2 min, and the supernatant containing the nucleoplasm was retained. The chromatin pellet was gently washed with 100 μL 1x MNase Buffer (NEB). Subsequently, 100 μL of pre-warmed Chromatin Digest Buffer (1x MNase Buffer, 0.2 mg/mL bovine serum albumin, 50 U/ μL MNase, 100 mM NaCl) was added and incubated at 37°C with agitation for 90 seconds. The digestion was terminated by adding ethylene glycol-bis(β-aminoethyl ether)-N,N,N′,N′-tetraacetic acid (EGTA) to a final concentration of 250 mM. The digested sample was centrifuged at 13,000 g, 4°C for 5 min and chromatin-associated proteins were recovered from the supernatant. Proteins in relevant fractions were precipitated using trichloroacetic acid and analysed by western blotting (**Supplementary Table S4**).

## Supporting information

Memet_Supplementary Data 1

Memet_Supplementary Data 2

Memet_Supplementary Data 3

Memet_Supplementary Data 4

Memet_Supplementary Data 5

Memet_Supplementary Data 6

Memet_Supplementary Data 7

## DATA AVAILABILITY

Deep sequencing data presented in this article are available in the Gene Expression Omnibus (GEO) database [http://www.ncbi.nlm.nih.gov/geo/] under the following accession Codes: GSE336932 and GSE336020.

The mass spectrometry proteomics data have been deposited to the ProteomeXchange Consortium via the MaSSIVE partner repository [https://massive.ucsd.edu/ProteoSAFe/static/massive] with the dataset identifier: MSV000102250.

Code is available via GitHub [https://github.com/Herzel-lab/GPATCHfamily].

## ACKNOWLEDGEMENTS

We thank Rebecca Rossen Falk and Soraya Beneddine for conducting preliminary experiments, Sabrina Stille for technical support and Jimena Davila Gallesio for help with preparation of figures. The authors would like to thank the HPC Service of FUB-IT, Freie Universität Berlin, and the Scientific Compute Cluster at GWDG, the joint data center of Max Planck Society for the Advancement of Science (MPG) and University of Göttingen for computing time. Mass spectrometry experiments were performed using the infrastructure of the UMG Core Facility Proteomics. This work was supported by the Deutsche Forschungsgemeinschaft; SFB1565, project number 469281184 to AMO (P02), HU (P04), MTB (P06), KEB (P12), CL (Z02) and LH (associated project).

## AUTHOR CONTRIBUTIONS

IM, NKa and AR perform experiments. CL and HU performed mass spectrometry. NKr and HN performed RiboMeth-seq. AMO analyzed chromatin association. LH performed bioinformatics analysis on IP-MS data, RIP- and RNA-seq data. KEB and MTB conceived and coordinated the study. LH, KEB and MTB wrote the manuscript and all authors were involved in reviewing/editing.

## DISCLOSURE OF POTENTIAL COMPETING INTERESTS

The authors declare no competing interests.

## SUPPLEMENTARY INFORMATION

**SUPPLEMENTARY DATA – see associated .xlxs sheets**.

**Supplementary Data 1. IP-MS analysis of human G-patch proteins**.

**Supplementary Data 2. IP-MS analysis of DHX15**. DHX15 and G-patch proteins recovered with DHX15 are marked in red.

**Supplementary Data 3. In vitro ATPase activity of G-patches and DHX15/DHX16 alone or together with G-patches**.

**Supplementary Data 4. RNA binding of G-patches and DHX15 together with G-patches**.

**Supplementary Data 5. RNA interactomes of nucleoplasmic G-patch proteins analyzed by RIP-seq**.

**Supplementary Data 6. Analysis of snRNA 2′-*O*-methylation in cells depleted of ZGPAT and/or DHX15**.

**Supplementary Data 7. Alternative splicing and differential expression of cells depleted of individual G-patch proteins**.

## SUPPLEMENTARY FIGURES AND LEGENDS

**Supplementary Figure S1.**
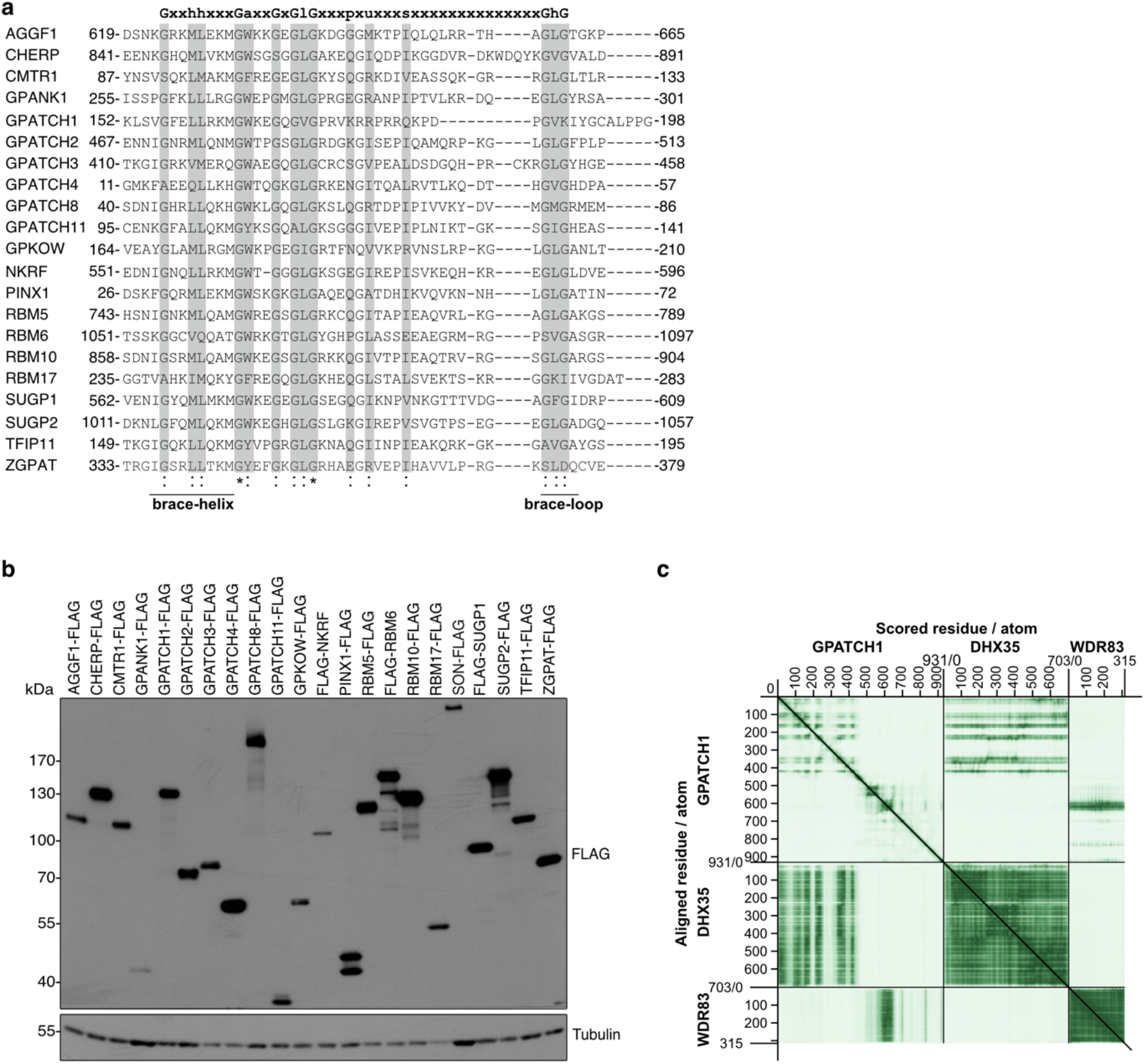
Alignment of G-patch sequence and expression of G-patch proteins in cells. **(a)** Alignment of the amino acid sequences of the G-patches of the 22 human G-patch protein according to UniProt. Amino acids of the G-patch consensus motif are highlighted in grey. Common amino acids are marked below the panel wherein * indicates fully conserved residues and : marks conservation between residues of similar properties. **(b)** Expression of FLAG-tagged versions of the indicated G-patch proteins was induced in stably transfected cell lines and whole cell extracts were analyzed by western blotting using antibodies against the FLAG tag and, as a loading control, tubulin. **(c)** Predicted aligned error (PAE) scores of the DHX35GPATCH1–WDR83 model (ipTM = 0.52pTM = 0.49) shown in Fig. 1h.

**Supplementary Figure S2.**
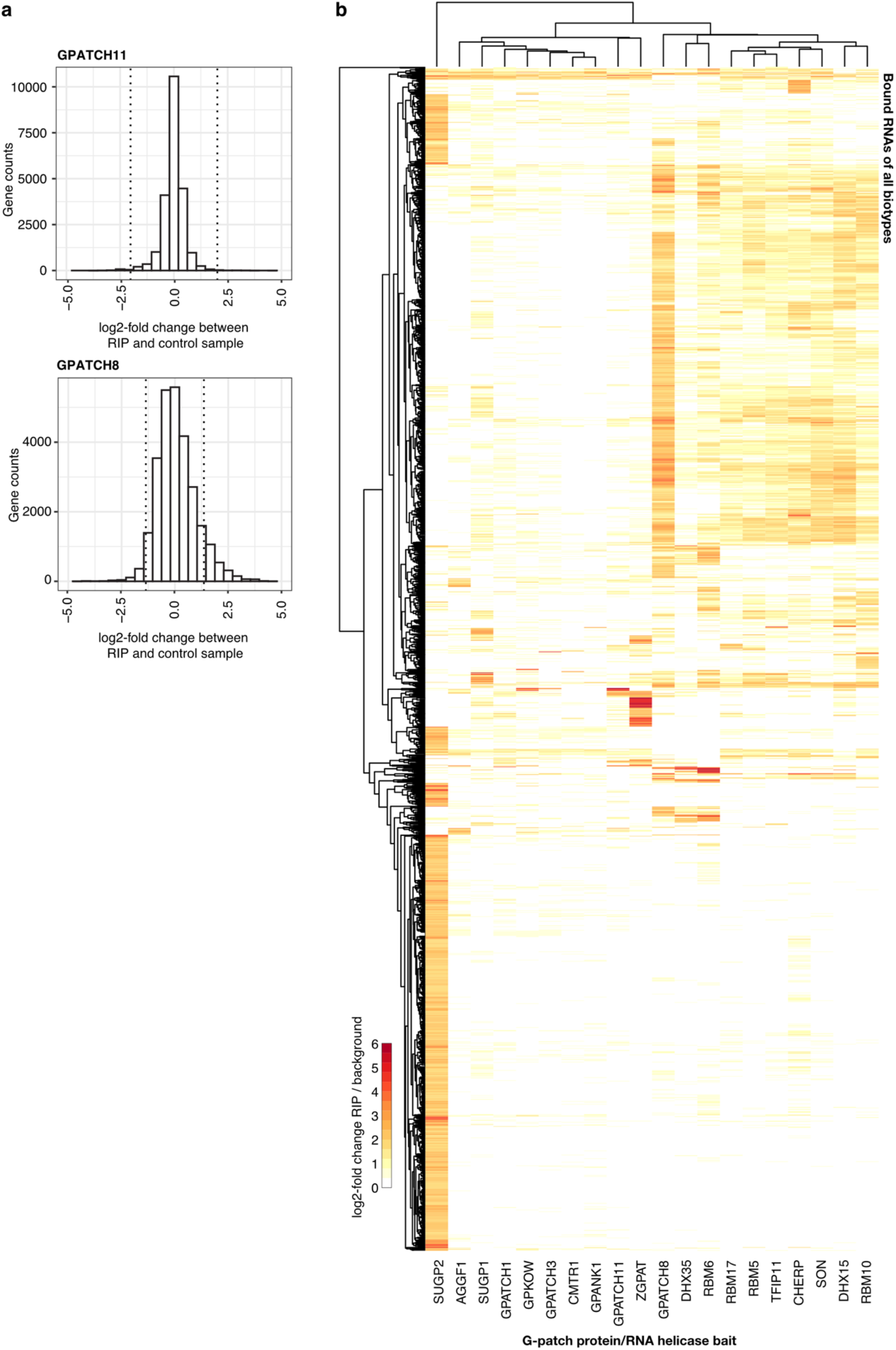
RNA interactomes of G-patch proteins and associated RNA helicases. **(a)** Distribution of log2-fold changes in RNA abundance in RIP-seq sample relative to FLAG control. GPATCH11 represents a distribution that led to the identification of few bound RNAs and for GPATCH8 hundreds of bound RNAs were identified. The dotted lines indicate the employed log2-fold change cutoff. **(b)** Heatmap showing hierarchal clustering of transcripts associated with the indicated G-patch proteins/RNA helicases.

**Supplementary Figure S3.**
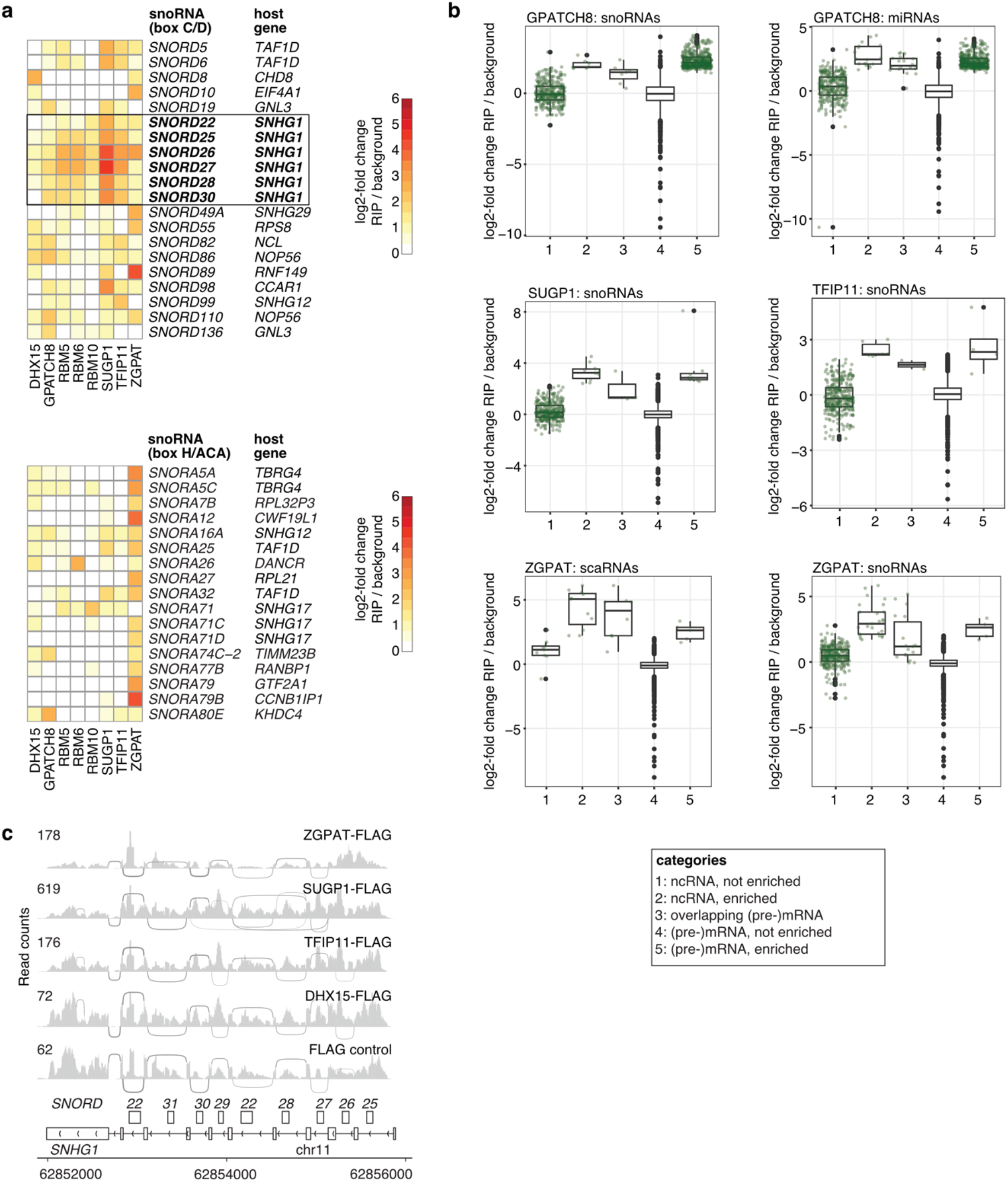
Association of G-patch proteins and DHX15 with ncRNAs and transcripts with intronic ncRNAs. **(a)** Heatmaps showing signal increase over background (FLAG control) for snoRNA genes in the RIP-seq datasets of the indicated proteins. The host genes within which each snoRNA is encoded is indicated; box and bold text highlight snoRNAs transcribed as part of the long non-coding snoRNA host gene SNHG1. **(b)** Boxplots showing distribution of ncRNA and (pre)mRNA signal over background for different RIP-seq datasets and type of ncRNA. Category 1, 4 and 5 serve as reference to compare how enriched certain ncRNAs / (pre)mRNAs are. Individual data points are labelled green for categories with less than 500 points, which means they were not plotted for category 4. **(c)** Read coverage for selected RIP-seq datasets and the FLAG control over the *SNHG1* gene with associated snoRNAs. Evidence of pre-mRNA splicing based on split reads is given by the bended lines, line thickness scales with number of split reads.

**Supplementary Figure S4.**
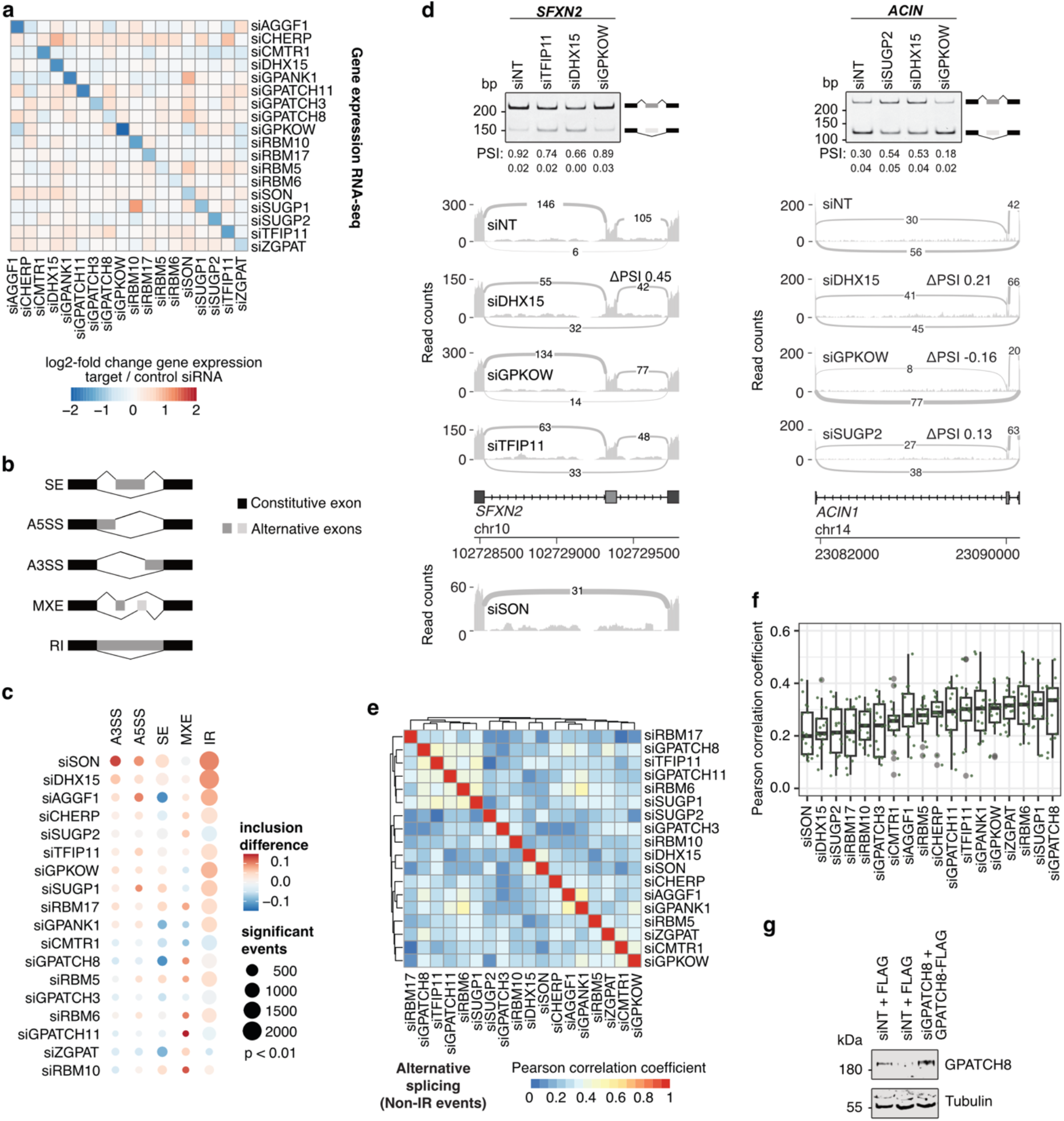
Differential expression and alternative splicing upon knock-down of individual G-patch proteins and DHX15. **(a)** Heatmap showing differential expression of *G-patch protein* and *DHX15* mRNAs upon siRNA-mediated depletion of the indicated G-patch proteins/RNA helicase. **(b)** Schematic views of different types of alternative splicing: skipped exon (SE), alternative 5’ splice site (A5SS), alternative 3’ splice site (A3SS), mutually exclusive exon (MXE), retained intron (RI). **(c)** Different extent and directionality of global alternative splicing changes for groups of G-patch proteins/ DHX15. Dotplot of median inclusion difference for significant AS events (p-value < 0.01) and barplot of gene counts with and without significant AS events. Each RNA-seq dataset consists of n=3 biological replicates. Dot size scales with number of events, complements Fig. 6b. **(d)** RT-PCR validation of two significant SE events and corresponding sashimi plots for probed knock-downs (above gene diagram) and knock-downs that were not assayed by RT-PCR (below gene diagram). Sashimi plots reflect the cumulative coverage of n=3 biological replicates per condition. siNT coverage reflects a representative pool of three siNT replicates out of nine. For RT-PCR experiments, n=2 experiments were performed and percent spliced in (PSI) is given ± standard error of mean. **(e)** Comparison of global ΔPSI values of non-IR events by Pearson correlation. Events were included that were significant in at least one knock-down condition. **(f)** Distribution of Pearson correlation coefficients shown in (e) for individual G-patch protein or DHX15 knock-down to all other knock-downs. **(g)** Stably transfected HEK293 cell lines were treated with non-target siRNAs (siNT) or those targeting GPATCH8 (siGP8) and expression of the FLAG tag or GPATCH8-FLAG was induced. Proteins were analyzed by western blotting using the indicated antibodies.

**Supplementary Figure S5.**
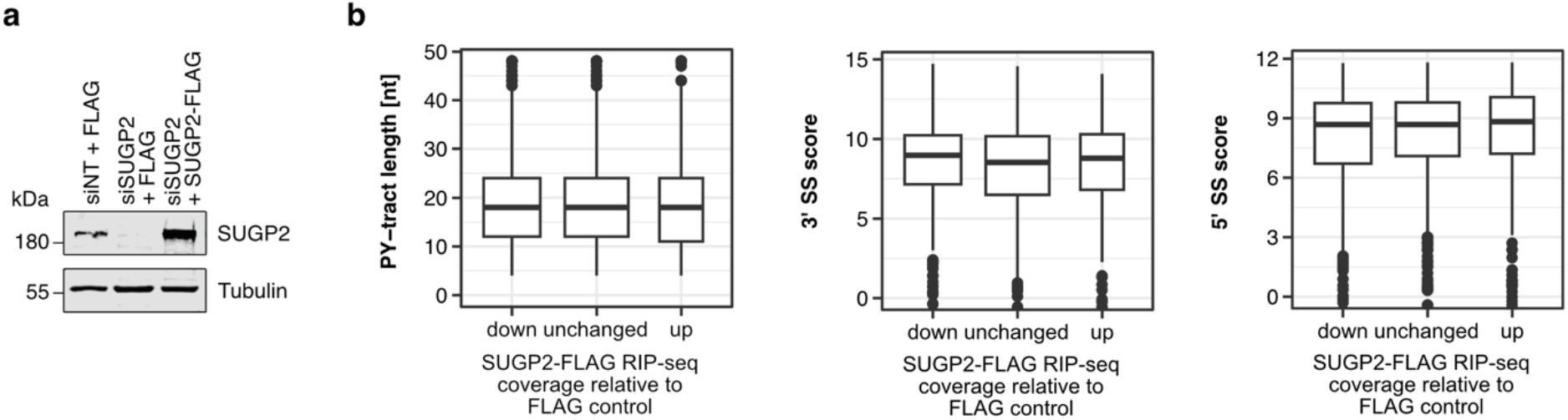
Splice site features of SUGP2-bound introns. **(a)** Stably transfected HEK293 cell lines were treated with non-target siRNAs (siNT) or those targeting SUGP2 (siSUGP2) and expression of the FLAG tag or SUGP2-FLAG was induced. Proteins were analyzed by western blotting using the indicated antibodies. **(b)** Distribution of pyrimidine (PY) tract lengths, 3’ and 5’ splice site (SS) scores grouped by signal in SUGP2 RIP-seq of the respective introns. No significances detected (p-Values > 0.05) by two-sided Wilcoxon rank sum test with Benjamini & Hochberg correction.

## SUPPLEMENTARY TABLES

**Supplementary Table S1:** Plasmids used and generated in this study.

| Plasmid name | CDS | Plasmid backbone | Source | Identifier |
| --- | --- | --- | --- | --- |
| AFFG1-His<sub>6</sub>-2xFLAG | NM_018046 | pcDNA5/FRT/TO | This study | pMB1339 |
| CHERP-His<sub>6</sub>-2xFLAG | NM_006387 | pcDNA5/FRT/TO | This study | pMB1338 |
| CMTR1-His<sub>6</sub>-2xFLAG | NM_015050 | pcDNA5/FRT/TO | This study | pMB1302 |
| His<sub>6</sub>-2xFLAG-DHX15 | NM_001358 | pcDNA5/FRT/TO | <sup>1</sup> | pMB420 |
| DHX35-His<sub>6</sub>-2xFLAG | NM_021931 | pcDNA5/FRT/TO | This study | pMB1472 |
| DHX35<sub>L460E</sub>-His<sub>6</sub>-2xFLAG | NM_021931 | pcDNA5/FRT/TO | This study | pMB1807 |
| His<sub>6</sub>-2xFLAG | - | pcDNA5/FRT/TO | <sup>2</sup> | pMB187 |
| GPANK1-His<sub>6</sub>-2xFLAG | NM_001199237 | pcDNA5/FRT/TO | This study | pMB1326 |
| GPATCH1-His<sub>6</sub>-2xFLAG | NM_018025 | pcDNA5/FRT/TO | This study | pMB1327 |
| GPATCH1G156E-His<sub>6</sub>-2xFLAG | NM_018025 | pcDNA5/FRT/TO | This study | pMB1809 |
| GPATCH2-His<sub>6</sub>-2xFLAG | NM_018040 | pcDNA5/FRT/TO | This study | pMB1328 |
| GPATCH3-His<sub>6</sub>-2xFLAG | NM_022078 | pcDNA5/FRT/TO | This study | pMB1329 |
| GPATCH4-His<sub>6</sub>-2xFLAG | KJ902705 | pcDNA5/FRT/TO | <sup>3</sup> | pMB1370 |
| GPATCH8-His<sub>6</sub>-2xFLAG | NM_001002909 | pcDNA5/FRT/TO | This study | pMB1487 |
| GPATCH8<sub>si_ins</sub>-His<sub>6</sub>-2xFLAG | NM_001002909 | pcDNA5/FRT/TO | This study | pMB2183 |
| GPATCH11-His<sub>6</sub>-2xFLAG | AK294697 | pcDNA5/FRT/TO | This study | pMB1330 |
| GPKOW-His<sub>6</sub>-2xFLAG | NM_015698 | pcDNA5/FRT/TO | This study | pMB1332 |
| His<sub>6</sub>-2xFLAG-NKRF | NM_001173487 | pcDNA5/FRT/TO | <sup>1</sup> | pMB1036 |
| PINX1-His<sub>6</sub>-2xFLAG | NM_017884 | pcDNA5/FRT/TO | This study | pMB1258 |
| RBM5-His<sub>6</sub>-2xFLAG | NM_005778 | pcDNA5/FRT/TO | This study | pMB1340 |
| His<sub>6</sub>-2xFLAG-RBM6 | NM_005777 | pcDNA5/FRT/TO | This study | pMB1603 |
| RBM10-His<sub>6</sub>-2xFLAG | NM_005676 | pcDNA5/FRT/TO | This study | pMB1369 |
| RBM17-His<sub>6</sub>-2xFLAG | NM_001145547 | pcDNA5/FRT/TO | This study | pMB1334 |
| His<sub>6</sub>-2xFLAG-SUGP1 | NM_172231 | pcDNA5/FRT/TO | This study | pMB1372 |
| SUGP2-His<sub>6</sub>-2xFLAG | NM_001017392 | pcDNA5/FRT/TO | This study | pMB1337 |
| SUGP2<sub>si_ins</sub>-His<sub>6</sub>-2xFLAG | NM_001017392 | pcDNA5/FRT/TO | This study | pMB1835 |
| TFIP11-His<sub>6</sub>-2xFLAG | NM_001008697 | pcDNA5/FRT/TO | This study | pMB1335 |
| ZGPAT-His<sub>6</sub>-2xFLAG | NM_001195653 | pcDNA5/FRT/TO | This study | pMB1333 |
| ZZ-AGGF1_619-665-His<sub>7</sub> | NM_018046 | pQE-80 (A15) | This study | pMB1501 |
| ZZ-CHERP_841-891-His<sub>7</sub> | NM_006387 | pQE-80 (A15) | This study | pMB1506 |
| ZZ-CMTR1_87-133-His<sub>7</sub> | NM_015050 | pQE-80 (A15) | This study | pMB1505 |
| MBP-DHX15-His<sub>10</sub> | NM_001358 | pQE-80 (A102) | This study | pMB1102 |
| MBP-DHX16-His<sub>10</sub> | NM_003587 | pQE-80 (A102) | This study | pMB1481 |
| MBP-DHX35-His<sub>10</sub> | NM_021931 | pQE-80 (A102) | This study | pMB41479 |
| ZZ-GPANK1_255-301-His<sub>7</sub> | NM_001199237 | pQE-80 (A15) | This study | pMB1513 |
| ZZ-GPATCH1_152-198-His<sub>7</sub> | NM_018025 | pQE-80 (A15) | This study | pMB1502 |
| ZZ-GPATCH2_467-513-His<sub>7</sub> | NM_018040 | pQE-80 (A15) | This study | pMB1496 |
| ZZ-GPATCH3_410-458-His<sub>7</sub> | NM_022078 | pQE-80 (A15) | This study | pMB1516 |
| ZZ-GPATCH4_11-57-His<sub>7</sub> | KJ902705 | pQE-80 (A15) | This study | pMB1492 |
| ZZ-GPATCH8_40-86-His<sub>7</sub> | NM_001002909 | pQE-80 (A15) | This study | pMB1503 |
| ZZ-GPATCH11_69-115-His<sub>7</sub> | AK294697 | pQE-80 (A15) | This study | pMB1502 |
| ZZ-GPKOW_164-210-His<sub>7</sub> | NM_015698 | pQE-80 (A15) | This study | pMB1495 |
| ZZ-GPKOW-His<sub>7</sub> | NM_015698 | pQE-80 (A15) | This study | pMB1417 |
| ZZ-NKRF_551-596-His<sub>7</sub> | NM_001173487 | pQE-80 (A15) | This study | pMB1517 |
| ZZ-PINX1_26-72-His<sub>7</sub> | NM_017884 | pQE-80 (A15) | This study | pMB1489 |
| ZZ-RBM5_743-789-His<sub>7</sub> | NM_005778 | pQE-80 (A15) | This study | pMB1499 |
| ZZ-RBM6_1051-1097-His<sub>7</sub> | NM_005777 | pQE-80 (A15) | This study | pMB1497 |
| ZZ-RBM10_858-904-His<sub>7</sub> | NM_005676 | pQE-80 (A15) | This study | pMB1514 |
| ZZ-RBM17_235-283-His<sub>7</sub> | NM_001145547 | pQE-80 (A15) | This study | pMB1498 |
| ZZ-SUGP1_562-609-His<sub>7</sub> | NM_172231 | pQE-80 (A15) | This study | pMB1490 |
| ZZ-SUGP2_1011-1057-His<sub>7</sub> | NM_001017392 | pQE-80 (A15) | This study | pMB1504 |
| ZZ-TFIP11_149-195-His<sub>7</sub> | NM_001008697 | pQE-80 (A15) | This study | pMB1515 |
| ZZ-ZGPAT_333-379-His<sub>7</sub> | NM_001195653 | pQE-80 (A15) | This study | pMB1500 |
| pOG44 | Flp recombinase | pOG | Thermo Fisher Scientific | pMB145 |

**Supplementary Table S2:**
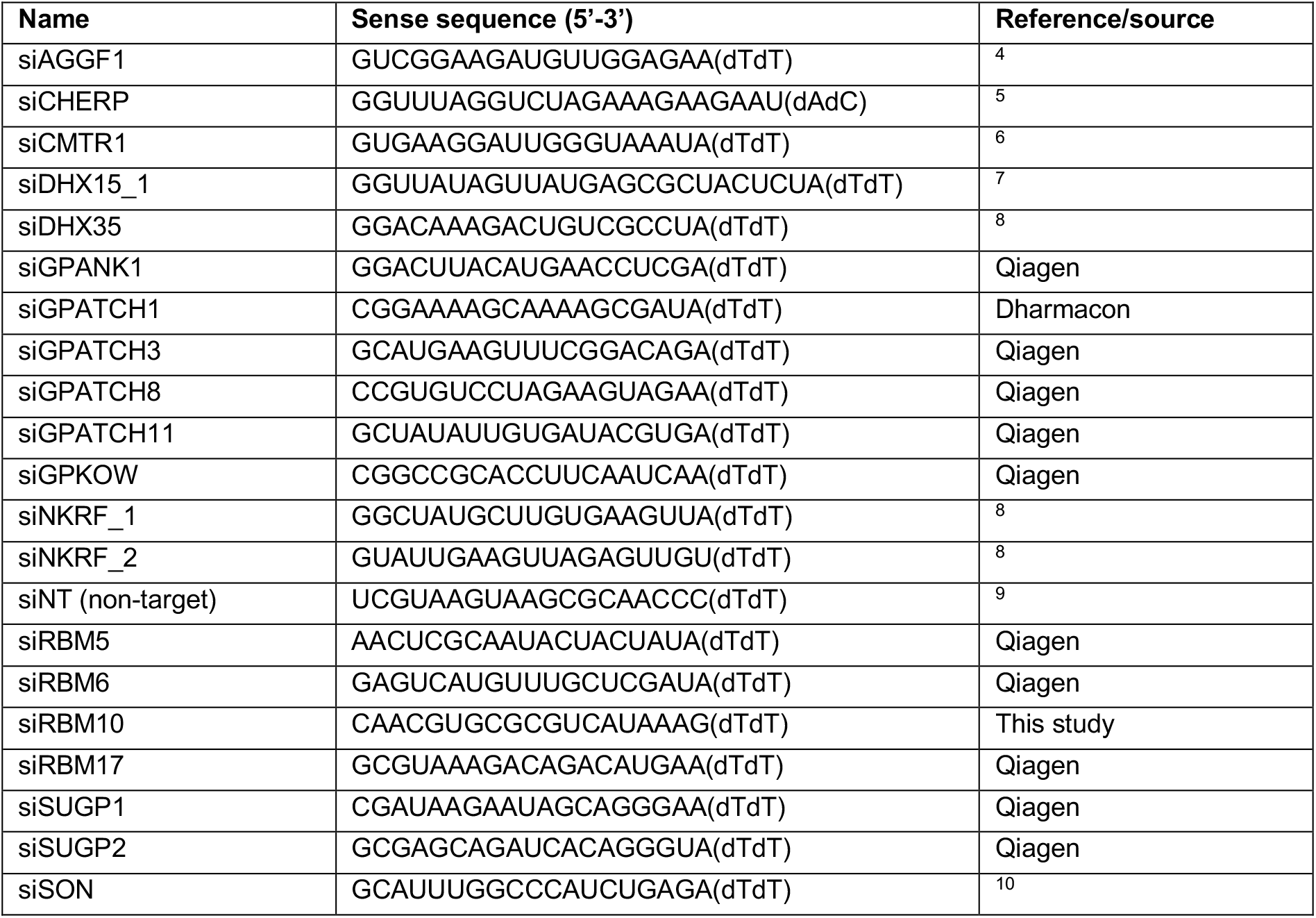
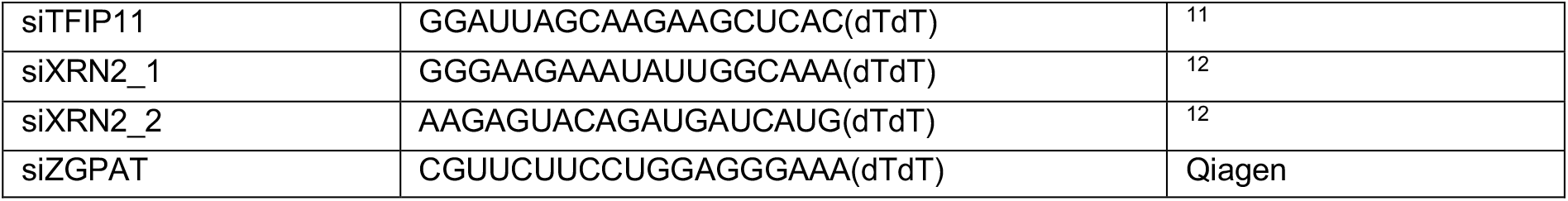
siRNAs used for RNAi-mediated depletion.

**Supplementary Table S3:**
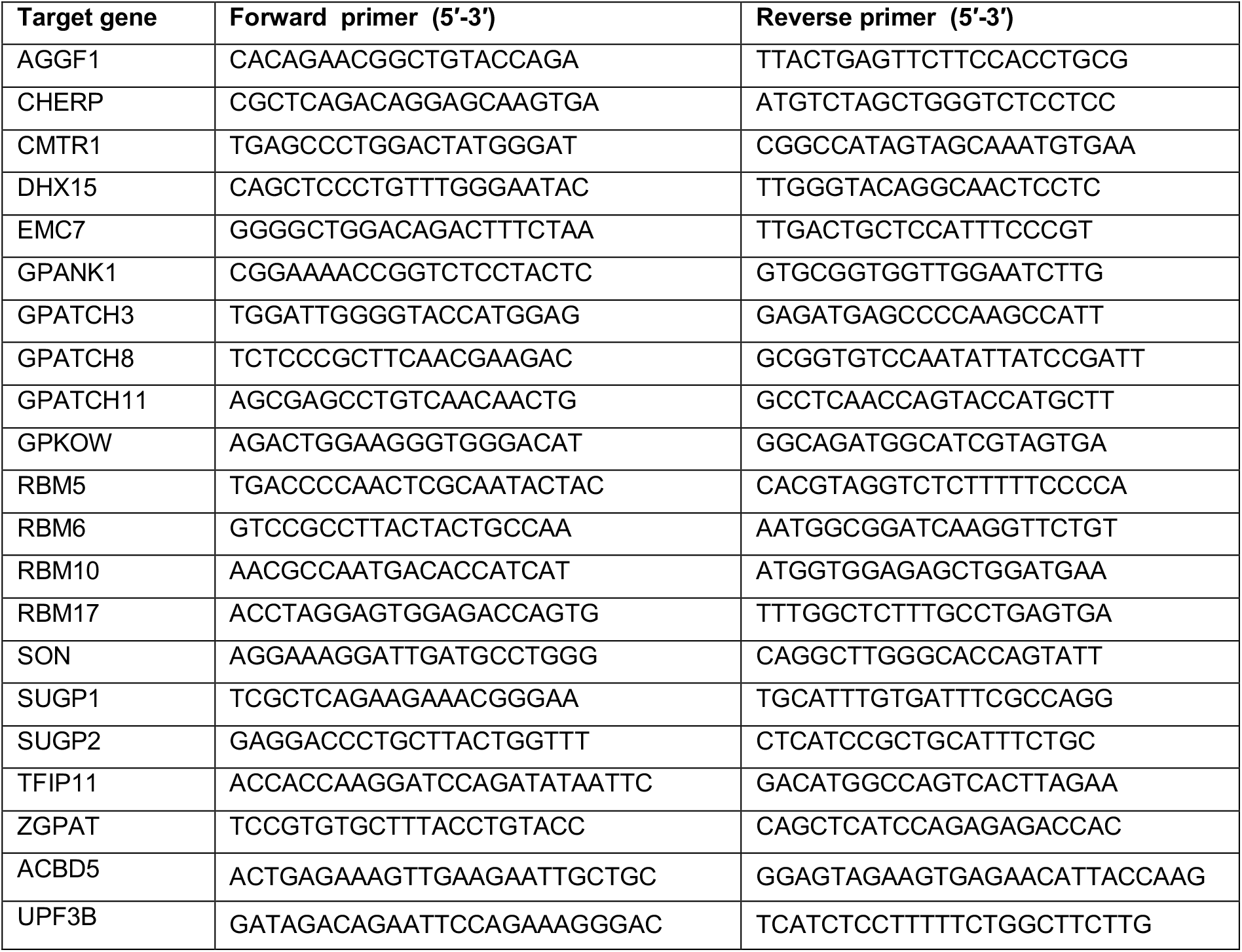
Oligonucleotide primers used for RT-(q)PCR primers.

**Supplementary Table S4:** Antibodies used in western blotting (WB) and immunofluorescence (IF).

| Name of antibody | Host | Dilution for WB | Dilution for IF | Source | # Cat. |
| --- | --- | --- | --- | --- | --- |
| DHX9 | Rabbit | 1:2000 | N/A | Bethyl | A300-854A |
| DHX15 | Rabbit | 1:5000 | N/A | Bethyl | A300-390A |
| DHX16 | Rabbit | 1:2000 | N/A | Bethyl | A301-537A |
| DDX21 | Rabbit | 1:2000 | N/A | Bethyl | A300-628A |
| DHX35 | Rabbit | 1:500 | N/A | Abcam | ab179442 |
| GPATCH1 | Rabbit | 1:250 | N/A | Abcam | ab130413 |
| WDR83 | Rabbit | 1:1000 | N/A | Proteintech | 27244-1-AP |
| Tubulin | Mouse | 1:10000 | N/A | Sigma-Aldrich | T6199 |
| FLAG M2 | Mouse | 1:5000 | 1:2000 | Sigma-Aldrich | F3165 |
| UTP14A | Rabbit | N/A | 1:100 | Proteintech | 11474-1-AP |
| ZGPAT | Rabbit | 1:2000 | N/A | Proteintech | 23203-1-AP |
| SUGP2 | Rabbit | 1:5000 | N/A | Proteintech | 12514-1-AP |
| Histone H3 | Rabbit | 1:5000 | N/A | Proteintech | 17168-1-AP |
| Anti-mouse - Alexa Fluor 488-conjugated | Goat | N/A |  | Jackson ImmunoResearch | 115-545-003 |
| Anti-mouse - Alexa Fluor 594-conjugated | Goat | N/A |  | Jackson ImmunoResearch | 115-585-003 |
| Anti-rabbit - Alexa Fluor 594-conjugated | Goat | N/A |  | Jackson ImmunoResearch | 111-585-003 |
| Anti-mouse - HRP-conjugated | Goat | 1:10000 |  | Jackson ImmunoResearch | 115-035-003 |
| Anti-rabbit - HRP-conjugated | Goat | 1:10000 |  | Jackson ImmunoResearch | 111-035-003 |
| Anti-rabbit - IRDye® 800CW | Donkey | 1:10000 | N/A | LI-COR | 926-32213 |
| Anti-rabbit - IRDye® 680LT | Donkey | 1:10000 | N/A | LI-COR | 926-68022 |

**Supplementary Table S5:**
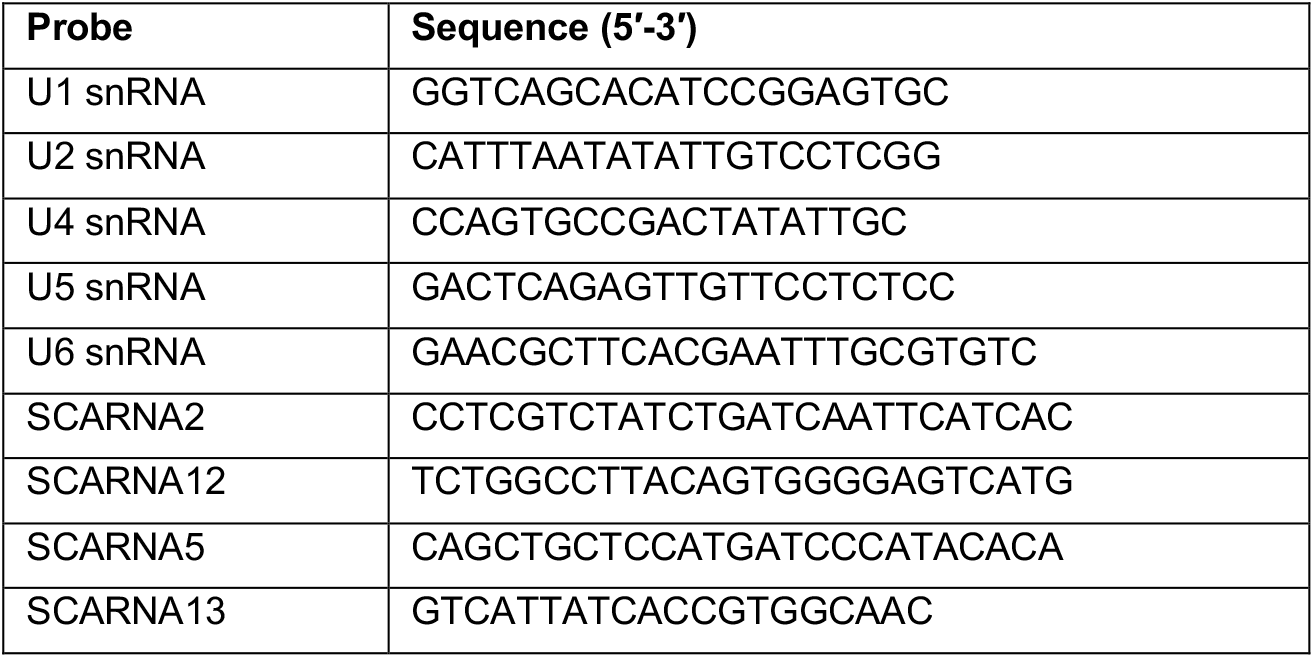
Oligonucleotides used as northern blot probes.

## SUPPLEMENTARY METHODS

### Multiple sequence alignment

The amino acid sequences of the G-patches of all human G-patch proteins as defined by UniProt were aligned using Clustral Omega^13^ using default parameters.

## Notes

### Competing Interest Statement

The authors have declared no competing interest.

